# Decoding the phenomenology of spontaneous thought using large language-model ratings on verbal retrospective free reports

**DOI:** 10.64898/2026.04.22.720079

**Authors:** Nicolás Bruno, Federico Cavanna, Federico Zamberlan, Tomás D’Amelio, Stephanie Muller, Laura Alethia de la Fuente, Jacobo Sitt, Antoni Valero Cabre, Mirta Villarreal, Enzo Tagliazucchi, Carla Pallavicini

## Abstract

Spontaneous thoughts constitute most of everyday inner experience, yet long-standing methodological challenges obscure a thorough exploration of their content and neurophysiological underpinnings. Traditional approaches relying on thought probes impose strict constraints on phenomenological reports, whereas online verbal reports disrupt the natural flow of experience while interfering neural signals with motor artifacts. Here, we designed and tested an alternative approach to assess the neural basis of spontaneous thoughts combining delayed verbal retrospective free reports (RFR) with automated phenomenological ratings generated by large language models (LLMs). Twenty-two participants performed an eyes-closed free-thinking task, providing reports that were evaluated along ten phenomenological dimensions by four state-of-the-art LLMs and a panel of human raters. Machine-learning models (ML) were then trained to decode LLM-derived ratings from EEG spectral, complexity, and connectivity features. Our analyses showed that inter-rater agreement among LLMs exceeded that of human raters whereas ML models achieved above-chance accuracy for the prediction of emotional valence. These findings provide support for the use of LLMs for a scalable phenomenological annotation of spontaneous thoughts and suggest that their affective dimensions can be decoded from concurrent EEG activity.

## 1 Introduction

Spontaneous thought, the continuous stream of mental content that unfolds in the absence of explicit task demands, occupies a substantial proportion of waking life. Estimates suggest that individuals spend approximately half of their time engaged in task-independent thoughts, commonly referred to as mind-wandering (Killingsworth & Gilbert, 2010; Smallwood et al., 2016). Accumulating evidence suggests that this activity may serve adaptive functions, including episodic future simulation, emotional regulation, autobiographical consolidation, and creative problem-solving (Andrews-Hanna et al., 2017; Corbani et al., 2025; Mildner & Tamir, 2024; Smallwood, 2013). The contents of spontaneous thought vary along several dimensions, such as temporal orientation (past, present, or future), emotional valence, self-relevance, sensory modality (verbal versus imagery), deliberateness, and vividness (Andrews-Hanna et al., 2013, 2022; Kirberg et al., 2025; Stawarczyk et al., 2011, 2013). Experience sampling studies have further shown that distinct patterns of phenomenological features cluster into separable cognitive states, such as goal-directed planning, ruminative self-reflection, and perceptual engagement (Coppola et al., 2024; Karapanagiotidis et al., 2017; Konu et al., 2021).

Understanding the neural basis of spontaneous thought is relevant for both fundamental and clinically applied neuroscience, as alterations in this process have been associated with psychiatric conditions such as depression, anxiety, and attention deficit hyperactivity disorder (ADHD) (Raffaelli et al., 2025; Seli et al., 2018; Smallwood et al., 2007). Human neuroimaging studies have consistently linked mind-wandering to activity within the default mode network (DMN), a set of midline and lateral cortical regions that exhibit preferential activation during internally oriented cognition (Andrews-Hanna et al., 2014; Christoff et al., 2016; Raichle et al., 2001). EEG and MEG, offering complementary temporal resolution, have further linked mind-wandering propensity to modulations in lower-frequency spectral power (Braboszcz & Delorme, 2011; Kam et al., 2022; Kawashima et al., 2023) and connectivity markers (Baird et al., 2014; Ibagon et al., 2023). However, most of these studies relied on thought-probe paradigms that constrain reports to predefined categories, leaving open the question of how the full phenomenological content of spontaneous thought maps onto neural activity.

Current approaches to capture the phenomenology of spontaneous thought can be divided into different categories depending on the constraints imposed on participants’ reports. At one end of this spectrum, *thought-probe* (TP) methods limit spontaneous thought reports to a predefined set of response options (Martinon et al., 2019; Smallwood et al., 2011). Despite its high temporal specificity, this approach could result in a potentially biased, phenomenological characterization of spontaneous thought by restricting participants’s choice to a list of predetermined dimensions (Andrews-Hanna et al., 2022; Stawarczyk et al., 2013). At the other end, *think-aloud paradigms* (TAP) involve the ‘live’ verbal narration of the stream of inner thoughts (Li et al., 2022; Raffaelli et al., 2021). While this approach allows for more nuanced and comprehensive reports, it may interfere with the flow of spontaneous cognition, potentially transforming freely unfolding thought into a qualitatively different, self-observed process (Fox et al., 2011; Gilles, Panneels, et al., 2025). Furthermore, speech-related muscle artifacts can severely compromise the quality of concurrently recorded EEG/MEG signals (Iwata et al., 2024; Van Calster et al., 2017).

Given these constraints, the current study proposes an alternative methodology designed to capture the phenomenological detail of spontaneous thought while minimizing interference from continuous vocal output, dubbed retrospective free report (RFR) methodology. This approach allows participants to think freely during a defined period, after which they are prompted to verbally describe their thoughts aloud with minimal constraints (Barzykowski et al., 2024). In this way, the proposed design aims to combine the strengths of TP and TAP paradigms: the subsequent verbal report preserves the open-ended, participant-driven phenomenological detail that questionnaires and forced-choice probes may fail to capture while the recording period remains free from speech-related artifacts preserving EEG/MEG signal quality.

While verbal reports provide more nuanced means of capturing the complex phenomenology of spontaneous thought, their analysis presents distinct methodological challenges. Compared with the numerical ratings obtained in more constrained designs, such as TP, verbal reports are less structured and therefore more difficult to quantify. Asking participants to reliably rate their experiences across multiple dimensions, as in standard experience sampling protocols (Andrews-Hanna et al., 2022; Gilles, Panneels, et al., 2025; Stawarczyk et al., 2014), becomes increasingly challenging as the number of trials and dimensions grows, extending task duration and introducing fatigue-related response biases. Delegating this evaluation to external human raters can reduce participant burden, but shifts the cost to trained coders, for whom the process may become prohibitively time-consuming given the number of reports per participant, and requires careful assessment of inter-rater reliability (Hurlburt & Heavey, 2006; Petitmengin, 2006). Critically, both approaches rely on subjective qualitative judgment, which may introduce systematic biases: heterogeneous rating criteria in self-reports, limited generalizability of trained raters’ assessments to other settings, and potential carryover effects from prior evaluations. Together, these issues may reduce the external validity of such approaches.

Addressing these limitations, our strategy combined the RFR paradigm with automated ratings of verbal reports of spontaneous thoughts generated by four commercially available large language models (LLMs). Compared with traditional natural language processing (NLP) methods (Corbani et al., 2025; Mallett et al., 2023; Mildner & Tamir, 2024), LLMs offer greater flexibility with minimal supervision and have recently been applied to the phenomenological analysis of subjective experience reports (Bertolini et al., 2024; Martínez-Pernía et al., 2025; Watkins, 2024). Participants performed a free-thinking task eyes closed (30-second trials), followed by immediately providing RFRs describing the content of their thoughts (see Figure 1.A). Multiple LLM agents and human expert raters evaluated each report along ten phenomenological dimensions, including valence, arousal, temporal orientation, self-relevance, sensory modality, plausibility, source, vividness, goal-directedness, and spontaneity, derived from the Multidimensional Experience Sampling (MDES) (Konishi et al., 2017; Konu et al., 2021; Smallwood et al., 2016; H.-T. Wang et al., 2018) (see Figure 1.B). Finally, a machine-learning pipeline was implemented to decode each phenomenological dimension from resting-state EEG features, including spectral power, complexity, and connectivity markers (see Figure 1.C).

**Figure 1:**
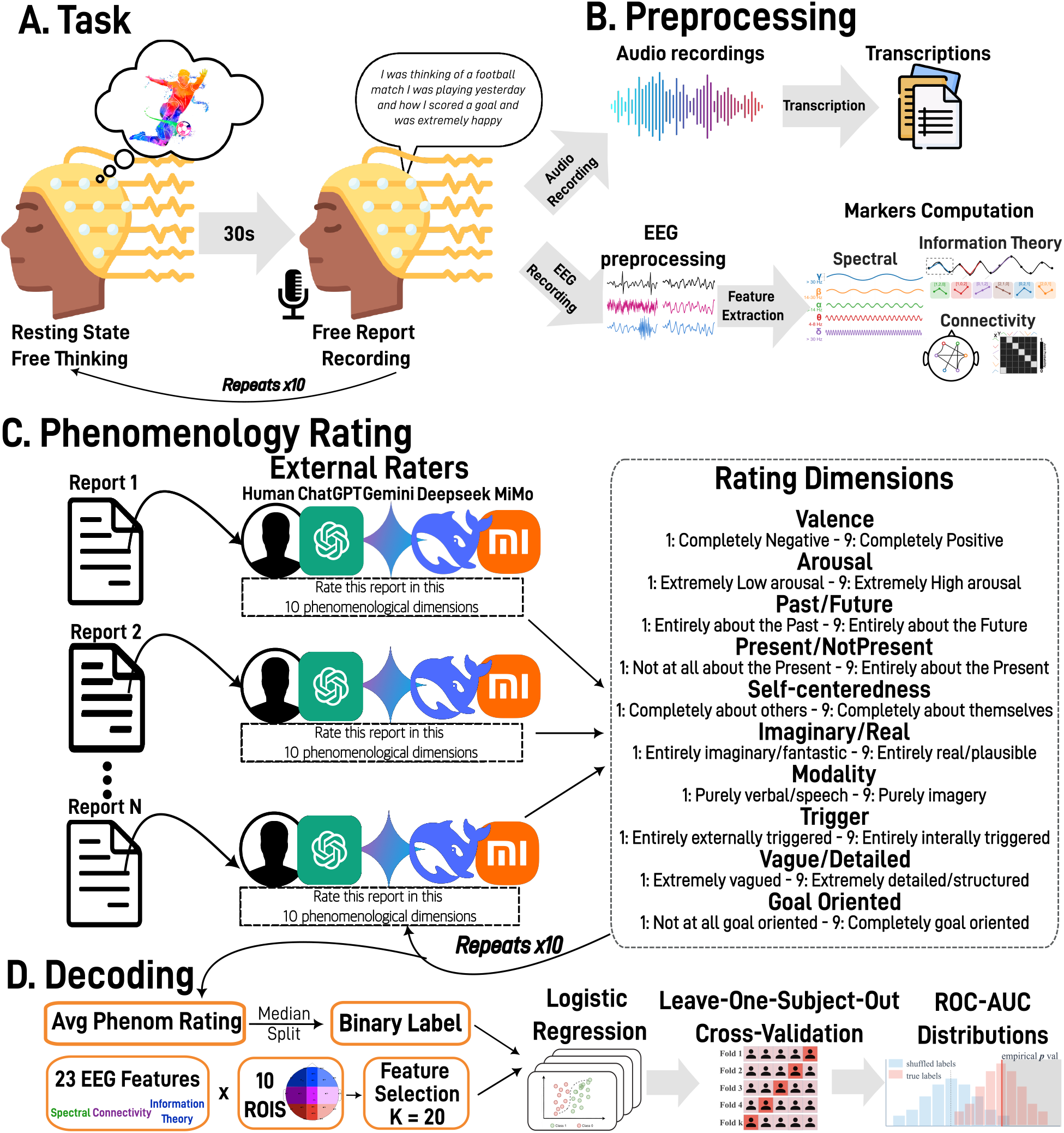
Experimental pipeline for EEG-based decoding of spontaneous thought phenomenology. **(A) Task:** Participants engaged in a 30-second eyes-closed free-thinking period (resting state) followed by retrospective verbal reports describing their thought content. This procedure was repeated for 10 trials per participant. **(B) Preprocessing:** Audio recordings were transcribed verbatim, while EEG data underwent standard preprocessing including filtering (1-90 Hz), artifact removal, and feature extraction. A comprehensive set of 23 EEG biomarkers was computed across spectral, information-theory, and connectivity domains, organized into 10 regions of interest (ROIs), comprising 9 regional ROIs based on anterior-posterior and laterality dimensions and one global scalp ROI. **(C) Phenomenology Rating:** Transcribed reports were evaluated along 10 phenomenological dimensions using both human raters and Large Language Models (LLMs). External human raters (n=52) provided ground truth ratings on 9-point Likert scales across dimensions including Valence, Arousal, Past/Future orientation, Present/NotPresent focus, Self-centeredness, Imaginary/Real content, Verbal/Imagery modality, Internal/External trigger, Vague/Detailed specificity, and Goal Orientation. Multiple state-of-the-art LLMs (ChatGPT-5.2, DeepSeek-V3, MiMo, Gemini-3-Flash) performed automated ratings using identical phenomenological definitions and rating scales. **(D) Decoding:** EEG features were used to predict binarized (within-subject median split) phenomenological dimensions through supervised machine learning. The pipeline employed Leave-One-Subject-Out cross-validation with feature selection (mRMR, k=20), elastic-net logistic regression classification, and SMOTE balancing. Model performance was assessed using ROC-AUC with permutation testing for statistical validation. SHAP values quantified feature importance to identify the most discriminative neurophysiological markers for each phenomenological dimension.

The current study aims to develop and evaluate a combined approach that integrates RFR paradigm, automated LLM-based phenomenological rating, and machine-learning EEG decoding as an ecologically feasible means of probing the neural correlates of spontaneous thought content. To this end, three empirical questions are addressed: whether LLMs can achieve phenomenological agreement with human raters comparable to inter-human consistency; whether automated annotation captures the phenomenological structure of spontaneous thought under the RFR paradigm; and whether LLM-derived ratings carry sufficient biological signal to be decoded from concurrent resting-state EEG.

## 2 Methods

### 2.1 Participants

A cohort of 22 participants (15 females, 36.1±13.1 years, range 19–59 years) took part in the experimental evaluation of our new strategy to collect and assess the content while decoding neural EEG markers associated to its main features. All participants were native Spanish speakers. This study was conducted in accordance with the Declaration of Helsinki and approved by the Research Ethics Committee at FLENI (Buenos Aires, Argentina). All participants gave written informed consent and received no financial compensation for their participation.

### 2.2 Materials

Electrophysiological data were recorded using a 24-channel wireless EEG SMARTING system (mBrainTrain LLC, Belgrade, Serbia; https://www.mbraintrain.com/) coupled with an elastic electrode cap (EASYCAP GmbH, Inning, Germany). Twenty-four Ag/AgCl electrodes were positioned according to the international 10–20 system at the following locations: Fp1, Fp2, Fz, F7, F8, FC1, FC2, Cz, C3, C4, T7, T8, CPz, CP1, CP2, CP5, CP6, TP9, TP10, Pz, P3, P4, O1, and O2. The reference and ground electrodes were placed at FCz and AFz, respectively. Data acquisition was performed with a 24-bit resolution DC amplifier (sampling rate: 500 Hz) securely attached to the electrode cap. EEG signals were digitized and transmitted via Bluetooth to a recording notebook for subsequent processing.

### 2.3 Procedure

Participants were seated in a comfortable chair within a quiet and dark room. Initially, a 1-minute baseline resting-state EEG recording (eyes-closed) was acquired. Following this baseline, the RFR paradigm was implemented. During each experimental trial, participants were instructed to remain relaxed, maintain their eyes closed and a neutral facial and body expression for 30 seconds (Figure 1A). Within this interval, participants were asked to engage in free-thinking, allowing their thoughts to unfold freely spontaneously in absence of any specific instruction or constraint on the investigator’s side. Critically, no speech occurred during this period, hence EEG data were continuously recorded with minimal muscle artifact. Immediately after each 30-second free-thinking interval, participants were prompted to provide a retrospective verbal report, describing the content of their thoughts aloud in Spanish with as much detail as possible. These reports were collected using a digital audio recorder. This trial structure was repeated 10 times per participant, yielding a total dataset of 220 retrospective verbal reports. Each complete experimental session lasted approximately 30 minutes.

### 2.4 Phenomenological Analysis

Audio recordings of the retrospective verbal reports were manually transcribed verbatim (Figure 1B). Subsequently, to characterize the phenomenological content of these reports, an adapted version of the MDES (Konishi et al., 2017; Konu et al., 2021; Smallwood et al., 2016; H.-T. Wang et al., 2018) was employed for external evaluation by both human raters and LLMs (Figure 1C). The evaluated dimensions included: Valence, Arousal, Past/Future, Present/NotPresent, Self-centeredness, Imaginary/Real, Verbal/Imagery Modality, Internally/Externally Triggered, Vague/Detailed, and Functionality/Goal Oriented. All dimensions were rated on a 9-point Likert scale. To establish the reliability of the automated rating pipeline, LLMs evaluations were systematically benchmarked against human raters.

#### 2.4.1 Human Raters

To establish a ground truth for the phenomenological dimensions, the entire set of 220 reports was divided into 16 non-overlapping randomized subsets (approximately 16 reports per subset, with no report appearing in more than one subset). These subsets were distributed via an online survey platform (Google Forms) that randomly assigned each human rater to one of the 16 subsets. A total of 52 independent anonymous volunteers participated in the rating task. Participants were permitted to rate multiple subsets if desired; on average, each participant rated 35.9 reports, resulting in approximately 8.11 ratings per report (range: 5–17).

#### 2.4.2 LLM automatic phenomenology analysis

The computational analysis was implemented in Python, utilizing the OpenRouter API (https://openrouter.ai/) to interface with the selected LLMs. A critical methodological feature of this approach was the enforcement of strict independence between evaluations. Unlike human raters, who assessed a sequence of reports and were potentially susceptible to context or order effects, the models processed each verbal report in total isolation. Specifically, each API call consisted of a single prompt containing one report, with no conversational history retained between interactions. This stateless design ensured that the automated ratings remained blinded and unbiased by prior inputs, eliminating potential carry-over effects inherent in sequential human annotation.

Diverse prompt strategies were evaluated to identify the optimal cost-effective zero-shot prompting strategy (Brown et al., 2020), wherein the models performed the evaluation relying exclusively on the provided instructions and their pre-trained knowledge, without the aid of few-shot examples (the complete prompt configuration is provided in Supplementary Materials Section 8.1.1; for details on the prompting comparison, see Section 8.1.2). Crucially, the evaluations were performed directly on the original Spanish transcripts without any intermediate translation step, relying on the models’ inherent multilingual capabilities to preserve the nuanced semantic content of the participants’ reports. To ensure strict methodological alignment with the human validation phase, the prompts were constructed to mirror the survey instruments. The models were provided with the exact phenomenological definitions and Likert scale anchors used by the human raters, ensuring that both evaluation modalities operated under an identical conceptual structure.

To maximize reproducibility, a fixed random seed and a consistent API provider were employed across all experimental runs. This standardization mitigates the stochastic nature of LLM outputs. Regarding sampling parameters, `top_p` was set to 1, indicating that the model considered the full probability distribution during token selection without nucleus sampling truncation. Additionally, `presence_penalty` and `frequency_penalty` were both set to 0. By disabling these penalties, models were prevented from artificially suppressing potential outputs due to repetition, ensuring that the generated ratings were driven exclusively by the prompt content rather than diversity-promoting mechanisms. Finally, the model’s temperature (a sampling parameter controlling output randomness, where *T* = 0 yields deterministic outputs and higher values increase stochastic diversity) was set to *T* = 1 to allow for greater diversity in generation, followed by a repetition of the rating procedure 10 independent times per report, and then averaging these 10 reports into one final value. The rationale for this stochastic aggregation strategy is that natural language often contains inherent semantic ambiguity; whereas a deterministic *T* = 0 setting forces the model into a single point estimate, sampling multiple times at *T* = 1 allows for the capture of a distribution of possible interpretations. This approach effectively marginalizes over the model’s uncertainty to yield a more robust and calibrated consensus rating that better reflects the nuances of subjective content. This strategy was formally compared against a deterministic zero-temperature (*T* = 0) single-shot baseline to ensure optimal performance (see Supplementary Methods 8.1.2 and Supplementary Figure S2).

These experiments were conducted utilizing two state-of-the-art, open-weights models: DeepSeek-V3 (DeepSeek-AI et al., 2025) and MiMo (Xiaomi et al., 2025). These models were specifically selected for their superior cost-performance ratio. As indicated by independent benchmarks (https://artificialanalysis.ai), they rank among the top-performing open-weights systems, offering minimal costs while maintaining high-level performance metrics comparable to more expensive alternatives. Also, the analysis was performed using high-performance proprietary models to evaluate the full potential of this methodology. Specifically, we employed widely recognized closed-weights systems, ChatGPT-5.2 (OpenAI, 2025) and Gemini-3-Flash (Google DeepMind, 2025), to assess the scalability and robustness of the results using the current state-of-the-art technology. A detailed breakdown of the costs associated with each model is provided in Supplementary Materials Table S1 (see Table S1 in Supplementary Materials). Notably, to further minimize token redundancy and optimize cost expenses, prompt caching mechanisms were utilized for compatible models (e.g., ChatGPT, Gemini & Deepseek).

#### 2.4.3 Benchmarking Phenomenological Ratings

For each verbal report and phenomenological dimension, a 20% trimmed mean rating across all human raters was computed and designated as the *human consensus score*. This aggregation method was employed to establish a robust measure of human consensus, minimizing the influence of outlier assessments. This aggregated value served as the ground truth for all subsequent model comparisons.

Agreement between each LLM and the human consensus was then assessed by computing Spearman’s rank correlation coefficient (*ρ*) between each model’s per-report ratings (each representing the average of 10 independent evaluations; see Supplementary Section 8.1.2) and the corresponding human consensus score across all 220 reports, separately for each phenomenological dimension, yielding a dimension-specific concordance measure per model. Pairwise agreement among LLMs — assessed independently of their correspondence with human ratings — is reported in Supplementary Section 8.2.2.

To contextualize the magnitude of these LLM-human correlations, a comparable human-level agreement baseline was established. For each individual human rater, Spearman’s *ρ* was computed between their ratings and the human consensus score (with that rater excluded from the consensus to avoid circularity). These individual coefficients were then averaged across all human raters, yielding a single summary value representing the typical level of agreement between an individual human rater and the group consensus. This human agreement baseline provided a direct reference against which to evaluate the LLM performance, where LLM-human correlations approaching or exceeding this value would indicate that the automated ratings achieved a level of concordance comparable to that observed among human raters.

To further corroborate the model’s performance on the emotional dimension, valence ratings were compared against established sentiment analysis tools, including a multilingual VADER implementation and a Spanish-adapted RoBERTa-based model fine-tuned on Twitter sentiment data (Loureiro et al., 2022). A detailed description of this comparative analysis is provided in Supplementary Results 8.2.3.

#### 2.4.4 Text Descriptors

To characterize the objective linguistic properties of the verbal reports, a set of text descriptors was extracted using the `textdescriptives` Python library (Hansen et al., 2023) integrated with spaCy’s large Spanish language pipeline (`es_core_news_lg`). The analysis pipeline incorporated multiple components: descriptive statistics, readability metrics, dependency distance measures, part-of-speech proportions, text quality indicators, and information theory metrics. These features were selected to capture complementary aspects of language production while minimizing redundancy. Report length was quantified via token count (number of words) and sentence count. Lexical diversity was assessed through Shannon entropy, reflecting the unpredictability of word sequences, where higher values indicate less stereotyped discourse. Syntactic complexity was indexed by the Flesch Reading Ease score, with lower values indicating denser, more complex text. Grammatical composition was characterized by the proportions of verbs, nouns, adjectives, and pronouns, capturing stylistic differences between action-oriented and object-focused descriptions. Finally, semantic coherence was computed at two levels: first-order coherence (average cosine similarity between adjacent sentences) and second-order coherence (similarity of each sentence to the document centroid), indexing local narrative flow and global topical consistency, respectively. Correlations between these objective text features and the LLM-derived phenomenological ratings were computed using Spearman’s *ρ*, with False Discovery Rate (FDR) correction for multiple comparisons.

### 2.5 EEG Data Processing and Feature Extraction

EEG data preprocessing was performed using the MNE-Python toolbox (Gramfort et al., 2013). The continuous EEG data were band-pass filtered between 1 and 90 Hz to remove slow drifts and high-frequency noise. A notch filter was applied at 50 Hz and its harmonics (50, 100, 150, 200 Hz) to eliminate power line interference. After removing by visual inspection noisy channels, the data was segmented into 2-second non-overlapping epochs. Automatic artifact rejection was applied using the AutoReject algorithm (random search method), which repairs or rejects bad epochs based on global and local peak-to-peak thresholds (Jas et al., 2017). Following this step, a visual inspection was conducted to manually identify and exclude any remaining noisy epochs or artifacts. Independent Component Analysis (ICA) was applied using the FastICA algorithm (Hyvarinen, 1999) to identify and remove ocular and muscle artifacts. Components reflecting artifacts were identified visually and subtracted from the data. Finally, bad channels were interpolated using spherical splines, and the data was re-referenced to the common average (CAR).

Following the framework proposed by Sitt et al. (2014) and Engemann et al. (2018), a comprehensive set of 23 EEG biomarkers was extracted (Figure 1B). These features are organized into three primary conceptual families (see Supplementary Table S2). First, spectral markers were derived from the Power Spectral Density (PSD) computation across standard frequency bands (*δ, θ, α, β*, and *γ*). Specific measures included both absolute and relative (normalized) band power, in addition to spectral summary metrics such as median frequency, spectral edge frequency, and spectral entropy. Second, information-theory markers, including Kolmogorov complexity and permutation entropy (PE), were calculated to quantify signal unpredictability. PE evaluates the diversity of local ordinal patterns, where higher values correspond to more complex and disordered signal dynamics, while lower values indicate more stereotyped activity. By adjusting the embedding delay (*τ*), PE assessed complexity across different temporal scales associated with each frequency band. Third, functional connectivity was estimated using weighted symbolic mutual information (wSMI). This non-linear approach transforms EEG signals into symbolic sequences to evaluate dependencies between channel pairs, effectively capturing long-range non-linear coupling while minimizing the influence of volume conduction.

Features were computed for each channel or channel pair and epoch. Crucially, to preserve spatial information, data were aggregated into 9 discrete ROIs (3×3) defined by anterior-posterior position (Frontal, Central, Posterior) and laterality (Left, Midline, Right), in addition to a global scalp ROI encompassing all channels (yielding a total of 10 ROIs; Figure 1D). The entire feature extraction pipeline was implemented using the Junifer EEG Python library (https://github.com/neurometers/junifer_eeg). To normalize the EEG features across participants, a baseline correction was performed: the mean feature vector obtained from a separate baseline resting-state recording was subtracted from the task-related feature vectors.

### 2.6 EEG Decoding Analysis

To assess whether the extracted EEG spectral features could predict the phenomenological dimensions derived from RFR reports, a supervised machine learning classification pipeline was implemented (Figure 1D). This analysis aimed to evaluate the extent to which neurophysiological features could predict reported subjective states.

#### Dimensions Binarization

The continuous phenomenological ratings were transformed into binary classes (“High” vs. “Low”) to facilitate robust classification. This transformation was achieved via a within-subject median split, ensuring maximum class balance. By defining the “High” class relative to each specific participant’s median rating rather than a global threshold, this approach prevented the classifier from exploiting stable individual differences (e.g., a participant who consistently rates Valence highly) as a proxy for the target variable of the decoding model. To ensure data quality, also a strict inclusion criterion based on class balance was implemented. Participants whose data exhibited extreme class imbalance (minority class containing less than 20% of samples) were excluded from the analysis for subsequent dimension-specific decoding, as such imbalance precludes reliable model training and evaluation.

#### Machine Learning Pipeline

To provide a non-artificial data augmentation strategy, models were trained and evaluated on each unaggregated 2-second epochs rather than report-level averages. An elastic-net Logistic Regression classifier was employed for all analyses. All features were standardized using within-subject z-score normalization to reduce inter-individual variability. Given the high dimensionality of the EEG feature space (23 features ×9 ROIs + Global Average) relative to the number of samples, the Minimum Redundancy Maximum Relevance (mRMR) algorithm was applied for feature selection. Within each cross-validation fold, the top 20 features (*k* = 20) were selected based on their relevance to the target class and minimum redundancy with other selected features. This selection step was performed strictly on the training data to avoid data leakage.

#### Model Performance and Validation

Model performance was evaluated using a Leave-One-Subject-Out (LOSO) cross-validation scheme. In each iteration, the model was trained on *N*−1 participants and tested on the participant held-out. This strategy strictly tests the model’s ability to generalize to unseen subjects (inter-subject decoding). Within each training fold, the Synthetic Minority Over-sampling Technique (SMOTE) was applied to synthetically augment the minority class, ensuring the classifier was trained on a balanced dataset.

To consolidate epoch-level predictions into a single classification per retrospective report, the predicted probabilities were averaged across all epochs belonging to that report. Final performance metrics, including the Area Under the ROC Curve (AUC), were calculated on these aggregated report-level predictions.

To determine the statistical significance of the classification performance, permutation testing was performed. For each model-dimension pair, a null distribution was generated by shuffling the target labels within subjects 1000 times and re-running the entire training and evaluation pipeline. This null distribution was compared to a distribution of 100 runs of the true model using different random seeds. The final empirical *p*-value was calculated as the proportion of permutation runs that achieved an AUC equal to or greater than the true model’s median AUC.

Moreover, on top of all the previously mentioned controls, and to further ensure that the model was not learning to predict participants instead of true target phenomenological labels, we conducted a validation analysis using synthetic data. Surrogate datasets were generated by preserving each participant’s covariance structure while destroying any relationship to the target labels. We verified that the full pipeline, when applied to this null data, maintained a Type 1 error rate below 0.05, meaning that significant classification results (where the “true” synthetic run outperformed the permuted runs) occurred in fewer than 5% of simulations. This step confirmed that the pipeline was robust against spurious associations driven by subject data clustering.

#### Feature Importance

To interpret the contribution of specific EEG features to the classification of phenomenological states, we computed SHAP (SHapley Additive exPlanations) values (Lundberg & Lee, 2017). SHAP values provide a unified measure of feature importance by quantifying the marginal contribution of each feature to the model’s output. For each fold of the LOSO cross-validation, absolute SHAP values were calculated and aggregated across all participants. To establish the significance of a feature’s contribution, the true mean absolute SHAP values were compared against two null baselines derived from the permutation testing procedure: (1) the mean SHAP value of that exact same feature across the permuted runs (“Shuffled (Same Feature)”), controlling for the feature’s inherent variance under the null hypothesis, and (2) the mean SHAP value of the feature occupying that same rank position across the permuted runs (“Shuffled (Same Rank)”), controlling for the expected magnitude of a top feature by pure random chance. This dual-baseline analysis allowed us to move beyond simple accuracy metrics and rigorously identify the electrophysiological signatures driving the successful decoding of subjective experience.

## 3 Results

### 3.1 Benchmarking LLM Phenomenological Ratings

All four LLMs surpassed the Human Rater Consistency baseline (Figure 2), achieving dimension-averaged correlations with the human consensus that exceeded the typical agreement among human raters (*ρ*_human_ = 0.63). Specifically, ChatGPT-5.2 obtained the highest mean *ρ* = 0.79, followed closely by Gemini-3-Flash (*ρ* = 0.78), DeepSeek-V3.2 (*ρ* = 0.75), and MiMo (*ρ* = 0.69). These results indicate that the automated LLM-based phenomenological ratings converged towards the human consensus more effectively than individual human judgments themselves.

**Figure 2:**
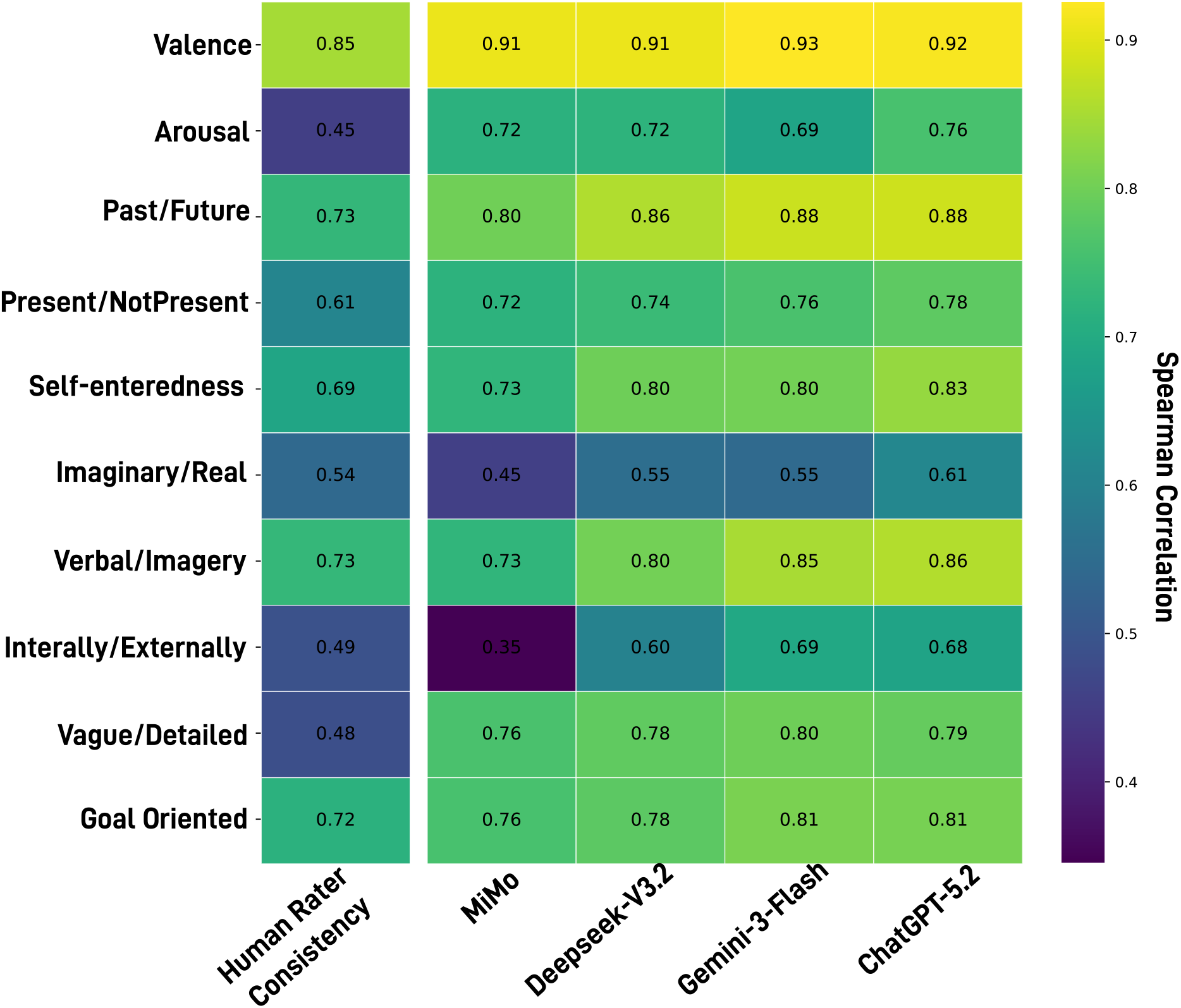
Agreement between LLMs and Human Consensus across Phenomenological Dimensions. Heatmap displaying Spearman’s rank correlations (*ρ*) between ratings from four LLMs (MiMo, DeepSeek-V3.2, Gemini-3-Flash, ChatGPT-5.2) and the human ground truth (average of *n* = 52 independent raters). The analysis covers ten dimensions: Valence, Arousal, Past/Future, Present/NotPresent, Self-centeredness, Imaginary/Real, Verbal/Imagery, Internally/Externally, Vague/Detailed, and Goal Oriented. The **Human Rater Consistency** column represents the reliability baseline, calculated as the average correlation between individual human raters and the group consensus. Warmer colors indicate stronger alignment. Results indicate that LLMs match or exceed human inter-rater reliability across most dimensions, particularly in Valence (*ρ >* 0.9), Past/Future and Verbal/Imagery Modality, suggesting they effectively capture the semantic structure of the reports.

At the dimension level, the strongest LLM-human agreement was observed for Valence (*ρ* = 0.91–0.93), Past/Future (*ρ* = 0.80–0.88), and Verbal/Imagery (*ρ* = 0.73–0.86). Self-centeredness (*ρ* = 0.73–0.83), Vague/Detailed (*ρ* = 0.76– 0.80), Goal Oriented (*ρ* = 0.76–0.81), and Present/NotPresent (*ρ* = 0.72–0.78) also exhibited consistent agreement across all models. In contrast, more modest correlations were obtained for Imaginary/Real (*ρ* = 0.45–0.61) and Internally/Externally (*ρ* = 0.35–0.69), dimensions for which Human Rater Consistency was also comparatively lower (*ρ*_human_ = 0.54 and 0.49, respectively). For Arousal, human raters exhibited poor consistency (*ρ*_human_ = 0.45), yet all four LLMs achieved substantially higher correlations with the human mean (*ρ* = 0.69–0.76). These results were corroborated by Intraclass Correlation Coefficient (ICC) analyses, which confirmed that LLMs achieved high rank-order correspondence as well as strong absolute agreement with the human consensus (see Supplementary Figure S3). Pairwise inter-model agreement was also assessed by computing Spearman correlations between all LLM pairs across each phenomenological dimension (see Supplementary Section 8.2.2 and Figure S4). Cross-model correlations were high for most dimensions (*ρ* = 0.76–0.96), except for Imaginary/Real and Internally/Externally, which exhibited greater variability across model pairs (*ρ* = 0.46–0.83).

To independently validate the models’ capacity to quantify emotional tone, LLM Valence ratings were compared against established sentiment analysis tools (VADER and RoBERTa). The generative LLMs achieved stronger agreement with the human consensus than both dedicated models and showed high inter-model agreement, correlating strongly with both VADER and RoBERTa (see Supplementary Analysis 8.2.3 and Supplementary Figure S5).

### 3.2 Phenomenological Profile of Retrospective Free Reports

Given that all four LLMs exceeded the Human Rater Consistency baseline and no substantial differences were observed across their rating distributions, ChatGPT-5.2 ratings were selected for all subsequent analyses as the best-performing model (*ρ* = 0.79).

Phenomenological ratings across the ten dimensions are displayed in Figure 3A. Regarding affective content, participants’ reports were predominantly rated as neutral to slightly positive in Valence (*M* = 5.50, *SD* = 1.65, *Mdn* = 6.0), whereas

**Figure 3:**
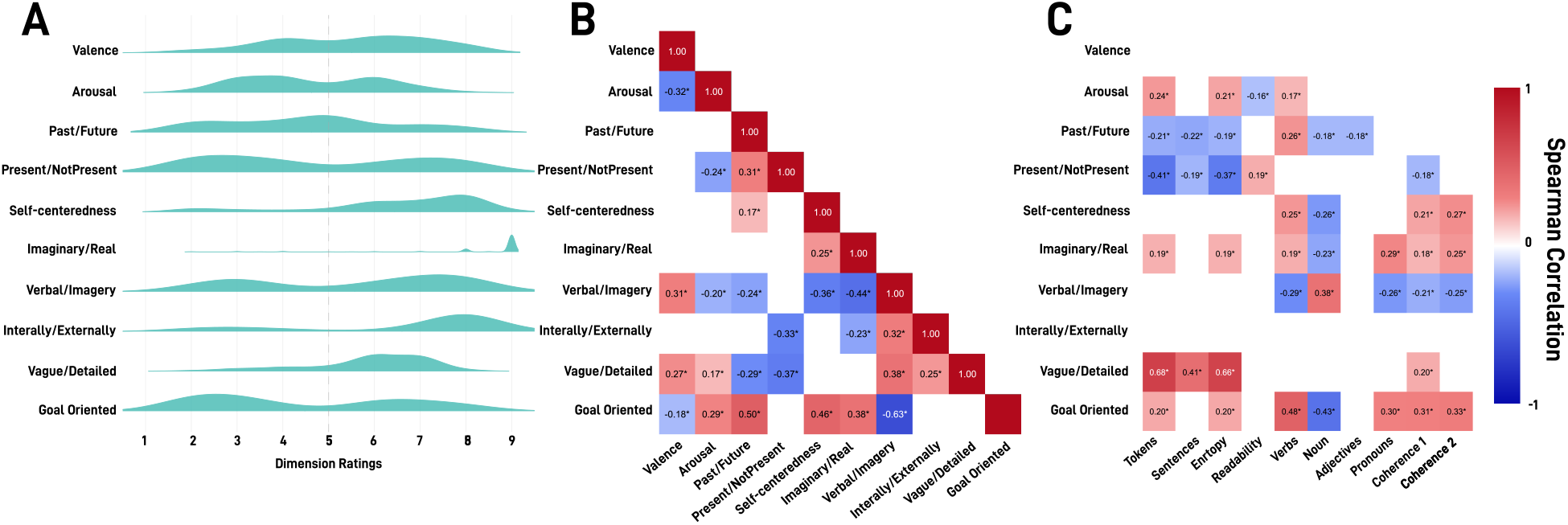
Phenomenological Profile of Resting-State Spontaneous Thought. **(A)** Score distributions across ten phenomenological dimensions rated by ChatGPT-5.2 (*n* = 220 reports), plots display the density of ratings on a 1–9 Likert scale. **(B)** Inter-dimensional correlations revealing the internal structure of phenomenological experience. Heatmap displays Spearman’s *ρ* coefficients between all dimension pairs (only FDR-corrected *p* < 0.05 shown). Red cells indicate positive associations (dimensions co-occur), blue cells indicate negative associations (dimensions are inversely related). Notable patterns include the negative Arousal-Valence link and Verbal/Imagery’s central role as a hub connecting multiple dimensions. **(C)** Correlations between phenomenological dimensions and objective text descriptors extracted via computational linguistics. Text features include report length (Tokens, Sentences), lexical diversity (Entropy), syntactic complexity (Readability), grammatical composition (Verbs, Nouns, Adjectives, Pronouns), and semantic coherence (first- and second-order). Only significant correlations are displayed (FDR-corrected *p* < 0.05). These associations validate the LLM ratings against objective linguistic markers.

Arousal (*M* = 4.63, *SD* = 1.46, *Mdn* = 4.0) exhibited a bimodal distribution, reflecting alternating states of low and higher activation. Temporal orientation was balanced between past and future (*M* = 4.63, *SD* = 1.85, *Mdn* = 5.0). The NotPresent/Present dimension (*M* = 4.80, *SD* = 2.24, *Mdn* = 4.0) also showed a bimodal distribution, indicating fluctuations between present-anchored and stimulus-independent thoughts. Self-centeredness ratings indicated consistent self-focus (*M* = 6.44, *SD* = 1.85, *Mdn* = 7.0), and Imaginary/Real ratings were notably high (*M* = 8.45, *SD* = 1.38, *Mdn* = 9.0), reflecting a strong predominance of realistic over imaginary content. Regarding representational modalities, Verbal/Imagery (*M* = 5.57, *SD* = 2.19, *Mdn* = 6.0) displayed a clear bimodal structure, suggesting thoughts tend to be either predominantly verbal or imagery-based. Vague/Detailed ratings were moderate (*M* = 5.87, *SD* = 1.31, *Mdn* = 6.0). Internally/Externally ratings (*M* = 6.60, *SD* = 2.22, *Mdn* = 8.0) suggested predominantly internally-driven thought, while Goal Oriented ratings (*M* = 4.48, *SD* = 2.18, *Mdn* = 4.0) were relatively low but with a bimodal tendency.

To examine the internal structure of phenomenological experience, Spearman correlations were computed between all dimension pairs (Figure 3B). Arousal exhibited a negative association with Valence (*ρ* = –0.32, *p*_FDR_ < 0.001), indicating that higher activation was linked to less positive affect. Verbal/Imagery emerged as a central hub, showing negative correlations with Self-centeredness (*ρ* = –0.36, *p*_FDR_ < 0.001), Imaginary/Real (*ρ* = –0.44, *p*_FDR_ < 0.001), and Goal Oriented (*ρ* = –0.63, *p*_FDR_ < 0.001), suggesting that imagery-rich thoughts tended to be less self-focused, less realistic, and less goal-directed. Conversely, Goal Oriented correlated positively with Self-centeredness (*ρ* = 0.46, *p*_FDR_ < 0.001), Imaginary/Real (*ρ* = 0.38, *p*_FDR_ < 0.001), and temporal orientation toward Past/Future (*ρ* = 0.50, *p*_FDR_ < 0.001), indicating that goal-oriented thoughts were more self-relevant, realistic, and temporally displaced. Internally/Externally correlated negatively with Present/NotPresent orientation (*ρ* = –0.33, *p*_FDR_ < 0.001), suggesting that internally-guided thoughts were less anchored in the present moment. Vague/Detailed showed positive associations with Verbal/Imagery (*ρ* = 0.38, *p*_FDR_ < 0.001) and Internally/Externally (*ρ* = 0.25, *p*_FDR_ < 0.001), and negative associations with temporal orientation (*ρ* = –0.30 to –0.37, all *p*_FDR_ < 0.001), indicating that more detailed reports were imagery-rich, internally-driven, and less temporally displaced.

To validate the phenomenological ratings against objective linguistic features, correlations were computed between LLM-derived dimensions and text descriptors extracted via computational linguistics (Figure 3C). Vague/Detailed exhibited the strongest associations, correlating positively with report length (Tokens: *ρ* = 0.68, *p*_FDR_ < 0.001) and lexical diversity (Entropy: *ρ* = 0.66, *p*_FDR_ < 0.001), confirming that subjectively detailed reports were objectively more elaborate. Goal Oriented correlated positively with verb proportion (*ρ* = 0.48, *p*_FDR_ < 0.001) and negatively with noun proportion (*ρ* = –0.43, *p*_FDR_ < 0.001), consistent with action-oriented, goal-directed content. Verbal/Imagery showed the inverse pattern, correlating positively with nouns (*ρ* = 0.38, *p*_FDR_ < 0.001) and negatively with verbs (*ρ* = –0.30, *p*_FDR_ < 0.001), reflecting the object-focused nature of imagery-based reports. Present orientation correlated negatively with report length (*ρ* = –0.41, *p*_FDR_ < 0.001) and entropy (*ρ* = –0.37, *p*_FDR_ < 0.001), suggesting that present-anchored thoughts yielded shorter, less elaborate descriptions.

### 3.3 EEG Decoding of Phenomenological Dimensions

To evaluate whether the subjective content of spontaneous thought could be predicted from underlying brain activity, a machine learning classification pipeline was employed to decode all ten phenomenological dimensions from resting-state EEG features. Classification of Valence achieved the highest decoding performance among all evaluated dimensions, demonstrating predictive capability significantly above chance level (AUC = 0.61, *p* = 0.006, FDR-corrected *p* = 0.043). Internally/Externally obtained the second highest AUC (0.57), though this did not reach statistical significance after correction (*p* = 0.11, FDR *p* = 0.34). Verbal/Imagery (AUC = 0.52, *p* = 0.32) and Goal Oriented (AUC = 0.52, *p* = 0.36) yielded near-chance performance. The remaining dimensions, Arousal (AUC = 0.49, *p* = 0.56), Self-centeredness (AUC = 0.48, *p* = 0.63), Vague/Detailed (AUC = 0.47, *p* = 0.74), Imaginary/Real (AUC = 0.46, *p* = 0.72), Present/NotPresent (AUC = 0.45, *p* = 0.90), and Past/Future (AUC = 0.42, *p* = 0.95), performed at or below chance level (all FDR-corrected *p* ≥ 0.67; Figure 4). Convergent evidence was obtained by applying the same decoding pipeline to discrete sentiment categories (Positive, Neutral, Negative) derived from ChatGPT-5.2, where Positive and Neutral sentiments were decoded above chance (AUC = 0.60 and 0.62, respectively; both FDR-corrected *p* = 0.015; see Supplementary Section 8.2.4). These results indicate that, within the current paradigm, the evaluated EEG markers carried discriminative information primarily for the emotional valence of spontaneous thoughts.

**Figure 4:**
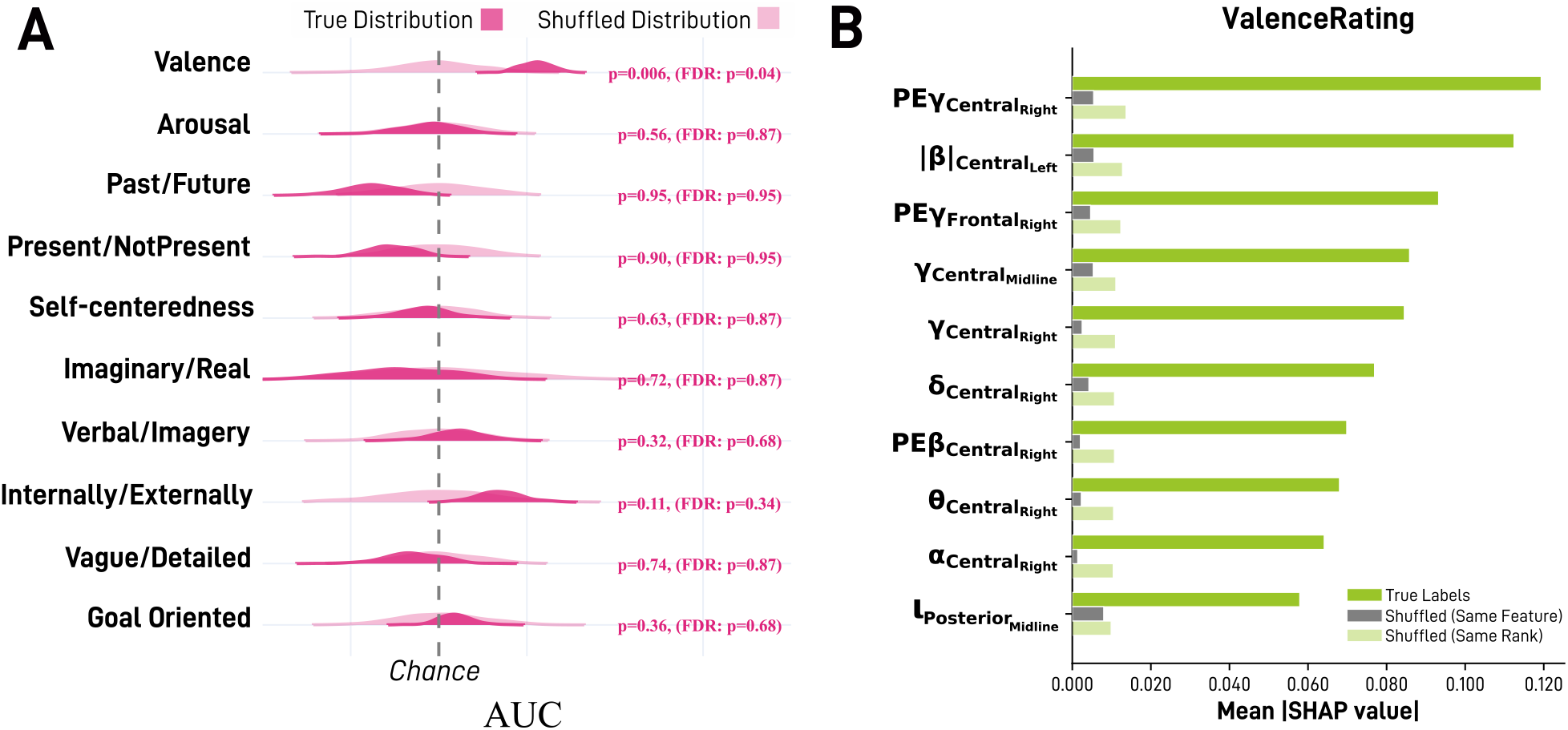
EEG Decoding and Neurophysiological Signature of Valence. **(A)** Violin plots displaying the final report-level Area Under the Curve (AUC) for the classification of all ten phenomenological dimensions, obtained after averaging epoch-level predictions. Dark pink distributions represent the true model AUC; light pink distributions represent the permutation-derived null. The dashed vertical line indicates theoretical chance level. Only Valence achieved statistically significant decoding performance (*p* = 0.006, FDR-corrected *p* = 0.043). **(B)** Feature importance profile for the Valence dimension. Bar plot displays the mean absolute SHAP values for its most discriminative EEG features. Solid green bars represent the true feature importance across all runs. Dark grey and light green bars denote the permutation-derived empirical baselines for the same feature and the same rank, respectively. Features substantially exceeding both baselines reflect robust neurophysiological signatures.

To further explore inter-individual variability in the predictability of mind-wandering experiences, a subject-level analysis was conducted (Figure 5). While the group-level cross-validation revealed that only Valence was consistently generalizable across the entire sample (Figure 4), per-subject decoding performance derived from each participant’s LOSO test fold demonstrated substantial heterogeneity. Across all participants, Valence remained the most consistently decodable construct at the individual level (reaching statistical significance in 45% of the subjects), followed by Goal Oriented (35%), and Verbal/Imagery (29%). The percentage of phenomenological dimensions that could be successfully decoded varied considerably among individuals. For instance, some participants exhibited highly predictable neurophysiological patterns, with up to 40% of the evaluated dimensions decoded significantly above chance, whereas others exhibited decoding performance that did not exceed chance levels for any dimension.

**Figure 5:**
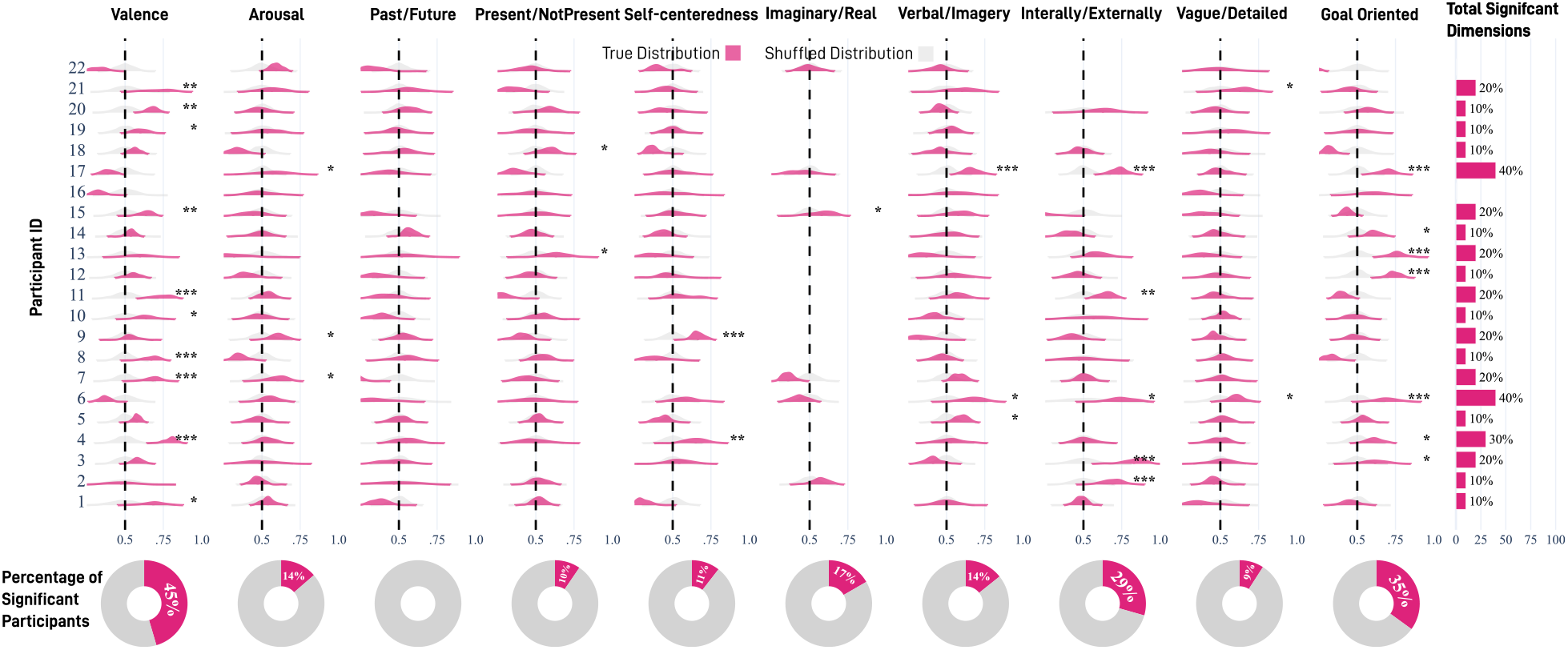
Subject-Level Decoding Performance of Phenomenological Dimensions. Violin plots display the cross-validated Area Under the Receiver Operating Characteristic Curve (ROC-AUC) distributions for individual subjects (rows) across the ten evaluated phenomenological dimensions (columns). Pink distributions indicate the model’s true predictive performance over multiple cross-validation iterations, whereas grey distributions represent the empirical null distribution derived from permutation testing with shuffled labels. The vertical dashed line in each cell denotes theoretical chance-level performance (AUC = 0.5). Statistical significance for each subject-dimension pair was determined by comparing the true performance against the permuted null distribution, denoted by asterisks (* *p <* 0.05, ** *p <* 0.01, *** *p <* 0.001). The rightmost panel (“Significant Dimensions per Participant”) presents a summary bar chart illustrating the percentage of phenomenological dimensions that were successfully decoded significantly above chance for each given subject, highlighting individual variability in the predictability of mind-wandering experiences. The bottom donut charts represent the percentage of subjects for which each specific dimension was decoded significantly above chance.

### 3.4 Neurophysiological Signature of Valence of Thought

To identify the specific electrophysiological features driving the decoding of spontaneous thought, we examined the SHAP values for the predictive models. Consistent with the cross-validated decoding performance, where only affective content achieved robust statistical significance, the feature importance analysis revealed a reliable neurophysiological signature primarily for the Valence dimension (Figure 4B).

For Valence, the predictive model relied on a distributed network of spectral and information-theoretic based markers over central and frontal regions. The most discriminative features consistently exceeded both permutation baselines, indicating statistically robust contributions. The predictive signature of emotional tone was primarily driven by three components: local signal complexity in high frequencies (gamma and beta permutation entropy: PE *γ*_Central, Right_, PE *γ*_Frontal, Right_, PE *β*_Central, Right_), regional spectral power across fast and slow rhythms (beta |*β*| _Central, Left_, gamma *γ*^Central, Midline^, *γ*^Central, Right^, delta *δ*^Central, Right^, theta *θ*^Central, Right^, and alpha *α*^Central, Right^), and posterior midline activity (Posterior_Midline_). Although other phenomenological dimensions lacked the group-level statistical significance and generalizable predictive power observed for Valence (Figure 4A), their full feature importance profiles can be found in Supplementary Figure S8.

## 4 Discussion

The current study aimed to develop and evaluate a combined approach integrating RFR paradigm, automated LLM-based phenomenological rating, and machine-learning EEG decoding as an ecologically feasible means of probing the neural correlates of spontaneous thought content. Three empirical questions guided this evaluation: whether LLMs could achieve phenomenological agreement comparable to inter-human consistency; whether automated annotation captured the phenomenological structure of spontaneous thought under the RFR paradigm; and whether LLM-derived ratings carried sufficient biological signal to be decoded from concurrent resting-state EEG. The present findings provide support for this approach, demonstrating that LLMs can reliably replicate human phenomenological judgments and that the resulting phenomenological dimensions map onto distinct electrophysiological signatures.

Investigating spontaneous thought presents two distinct methodological challenges. The first is a fundamental tension between capturing phenomenological richness and maintaining precise temporal indexing of neural activity. Thought probes, offer accurate localization at the cost of disrupting the natural flow of cognition (Martinon et al., 2019; Smallwood et al., 2011). The use of immediate retrospective verbal reports mitigates these artifacts by allowing thought to remain unconstrained, at the price of a more uncertain temporal indexing for these states. The second challenge is technical: extracting quantitative features from unstructured reports. Traditional qualitative approaches are labor-intensive and difficult to scale (Hurlburt & Heavey, 2006; Petitmengin, 2006), while previous computational attempts using basic embeddings or smaller language models often failed to capture complex psychological content (Bertolini et al., 2023, 2024; Li et al., 2022; Raffaelli et al., 2021).

Our findings suggest that the use of the most current-generation LLMs could overcome this technical limitation. All four models superseded Human Rater Consistency baseline (*ρ*_human_ = 0.63), with ChatGPT-5.2 achieving the highest concordance (*ρ* = 0.79). These outcomes extend recent work validating LLMs for descriptive phenomenological analysis (Martínez-Pernía et al., 2025; Shin & Luke, 2025) and sentiment annotation (Belal et al., 2023; Z. Wang et al., 2023). Unlike traditional NLP pipelines requiring domain-specific fine-tuning (Low et al., 2020; Sanz et al., 2022), the generative architecture of LLMs enables zero-shot scoring of complex psychological constructs defined through natural language instructions (Brown et al., 2020), positioning LLM-based rating as a scalable alternative that preserves dimensional richness while reducing the annotation costs that have historically limited sample sizes in experience sampling research (Martínez-Pernía et al., 2025). At the dimension level, agreement was highest for *Valence* (*ρ* = 0.91–0.93), *Past/Future* (*ρ* = 0.80–0.88), and *Verbal/Imagery* (*ρ* = 0.73–0.86), core phenomenological components whose manifestation in verbal reports tends to be linguistically transparent (Andrews-Hanna et al., 2013; Stawarczyk et al., 2013). Dimensions requiring greater inferential depth, such as *Internally/Externally Triggered* (*ρ* = 0.35–0.69) and *Imaginary/Real* (*ρ* = 0.45–0.61), showed more modest agreement, paralleling lower Human Rater Consistency for these constructs (*ρ*_human_ = 0.49 and 0.54), suggesting that the reduced agreement reflects inherent ambiguity in the constructs rather than a specific failure of the models (Engelbert & Carruthers, 2011; Hurlburt & Heavey, 2006). LLMs also achieved high consistency with the human consensus for *Arousal* (*ρ* = 0.69–0.76) despite markedly low inter-human agreement (*ρ*_human_ = 0.45), a stability further supported by complementary Intraclass Correlation analyses confirming strong absolute agreement (Supplementary Figure S3). These concordance values indicate that current LLMs can approximate inter-rater agreement for most phenomenological dimensions examined here, particularly those whose expression in verbal reports is linguistically transparent.

The observed phenomenological profile—characterized by high self-relevance (*M* = 6.44), plausibility (*M* = 8.45), and internal orientation (*M* = 6.60) with mildly positive valence (*M* = 5.50)—is consistent with the default mode of unconstrained cognition (Andrews-Hanna et al., 2014; Smallwood et al., 2016). Notably, several dimensions (Arousal, Present/NotPresent, Verbal/Imagery, and Goal Oriented) showed bimodal rather than uniform distributions, indicating that spontaneous thoughts tended to cluster at opposite ends of these scales. The pronounced plausibility and self-centeredness align with the established distinction between waking spontaneous thought and dreaming, indicating that mind-wandering remains constrained by reality monitoring and anchored in self-referential processing (Gross et al., 2021; Kirberg et al., 2025). Although goal orientation was relatively low (*M* = 4.48), its positive correlation with self-centeredness and temporal displacement suggests that a subset of thoughts retained an internally goal-directed quality, resembling the constructive/prospective component (Andrews-Hanna et al., 2013). The inter-dimensional correlation structure revealed two key organizational features. First, *Verbal/Imagery* operated as a central hub, negatively associated with self-centeredness, plausibility, and goal orientation. This aligns with findings that goal-directed, self-relevant thoughts are predominantly experienced as inner speech, whereas imagery-rich episodes tend to be less structured and less self-focused (Stawarczyk et al., 2013). Second, *Goal Oriented* thoughts clustered with self-centeredness, plausibility, and past/future orientation, mirroring the constructive/prospective factor consistently identified in experience sampling studies (Andrews-Hanna et al., 2013; Konu et al., 2021). Additionally, the negative arousal-valence association (*ρ* = −0.32) resembles the dysphoric rumination component linked to intrusive, negative mentation (Andrews-Hanna et al., 2013; Konu et al., 2021). Combined with convergent associations between phenomenological ratings and objective text descriptors (Öncel et al., 2025), the correlation structure observed here resembles the dimensional organization reported in probe-based studies (Andrews-Hanna et al., 2013; Konu et al., 2021), suggesting that the RFR paradigm with LLM annotation recovers comparable phenomenological features while providing a continuous characterization unconstrained by predefined probe categories.

Two features of the RFR paradigm, however, may have partially shaped this profile. Unlike TP methods—which more frequently sample fleeting, fragmentary, or even mind-blank states—the anticipation of a forthcoming report in a defined block may have promoted more narrative and self-relevant thought, consistent with evidence that awareness of an impending report shifts spontaneous cognition toward more structured and personally meaningful content (Gilles, Panneels, et al., 2025). Additionally, the retrospective nature of the report introduces a selective recall bias, whereby more coherent, positive, and self-referential episodes are better retained and therefore overrepresented (Gilles, Panneels, et al., 2025). Critically, both effects are substantially attenuated by the near-immediate delay of the RFR ( 30 seconds), which avoids the more severe encoding distortions associated with extended retrospective assessment, and by the absence of concurrent vocalization, which in TAP has been shown to alter the natural flow of spontaneous cognition (Fox et al., 2011; Garg et al., 2025; Gilles, Panneels, et al., 2025).

Among the ten phenomenological dimensions evaluated, only Valence was reliably decoded from resting-state EEG at the group level (AUC = 0.61, FDR-corrected *p* = 0.043), with convergent support from the successful classification of discrete sentiment categories (Positive AUC = 0.60, Neutral AUC = 0.62; both FDR-corrected *p* = 0.015). This selective result extends to EEG recordings prior fMRI work demonstrating that affective content is decodable from spontaneous thought (Kim et al., 2024; Tusche et al., 2014) and aligns with the broader observation that valence, as a fundamental organizing dimension of experience, tends to produce relatively stable and generalizable physiological signatures compared to more abstract phenomenological constructs (D’Amelio et al., 2025; Uusberg et al., 2013). Notably, the salience of valence during thought encoding and recall (Gilles, Panneels, et al., 2025) may further contribute to more consistent neural representations across individuals, facilitating group-level decoding. Also, several methodological factors may have further contributed to this asymmetry. Valence ratings were relatively well-distributed around the scale midpoint (*M* = 5.50), yielding balanced binary classes after median-split, whereas dimensions with highly skewed distributions, may have lost meaningful phenomenological contrast upon binarization, effectively collapsing physiologically distinct states into a single category. Additionally, while annotation noise from dimensions with lower LLM-human concordance could have obscured brain-behavior associations, this alone does not explain the full pattern, as well-agreed dimensions like Past/Future also failed to decode. The subject-level analysis revealed that the group-level null result for most dimensions may partly reflect inter-individual heterogeneity rather than a complete absence of neural correlates (Figure 5). Valence was the most consistently decodable dimension across participants (significant in 45% of subjects), but dimensions that failed at the group level, such as Goal Oriented (35%) and Verbal/Imagery (29%), were nevertheless decoded in a substantial proportion of individuals. This dissociation suggests that the neurophysiological correlates of these constructs may be highly idiosyncratic, consistent with evidence that neural representations of self-relevant content become increasingly person-specific (Anderson et al., 2025; Lux et al., 2022). The variability across participants points to a need for individualized modeling approaches in future work.

The RFR paradigm also carries intrinsic methodological limitations. Its most fundamental constraint is temporal: by collecting a single retrospective report for each 30-second interval, the approach cannot resolve rapid fluctuations within a trial. Given that brain states typically persist for only 6–8 seconds (Yu et al., 2025), a single report likely integrates several successive mental episodes. Brief states such as mind blanks or momentary attentional lapses, which TP methods may detect at the moment of occurrence, are systematically lost (Martinon et al., 2019). This reflects a broader tension inherent to experience-sampling methodology. Methods that prioritize temporal localization, such as instantaneous TPs or Temporal Experience Tracing (TET) (Lewis-Healey et al., 2025), restrict the phenomenological richness that can be accessed by interrupting the flow of thought. TAP methods achieve temporal continuity by externalizing thought as it unfolds, but the act of concurrent verbalization modifies what is experienced, selectively suppressing private or non-verbal content (Fox et al., 2011; Garg et al., 2025). The RFR preserves the natural trajectory of thought and allows for rich, unconstrained reports, but at the cost of temporal precision. Across these approaches, attempts to improve access to the content of ongoing thought appear inseparable from a corresponding cost, whether to temporal resolution, ecological validity, or measurement reactivity. A second limitation is that retrospective reports depend on memory. Thoughts that are more important, deliberate, or future-oriented are recalled preferentially (Gilles, D’Argembeau, & Stawarczyk, 2025), introducing a sampling bias in which the content reaching the report may not fully represent what was experienced. Furthermore, broad inter-individual differences in memory, metacognitive awareness, and verbal report quality may introduce systematic variance unrelated to the underlying mental states, a factor that could partly explain the marked variability in EEG decoding performance observed across participants.

While the present findings provide solid evidence for the proposed approach, several considerations may help guide future refinements. Regarding the LLM-based phenomenological assessment, the validation was conducted against external human raters rather than on self-reports, which means that the current pipeline captures inter-subjective consensus on what a report conveys rather than the participant’s own first-person experience. LLMs may therefore be best understood not as replacements for self-raters, but as tools that render external phenomenological annotation automatic, objective, and reproducible. Additionally, since human raters assessed sequential batches of reports and could have progressively fine-tuned their rating criteria as they accumulated exposure to more reports, whereas LLMs processed each report in complete isolation with no such calibration benefit, the LLM-human comparison may represent a conservative benchmark for automated rating performance. Future studies comparing LLM ratings against participants’ own self-assessments could help clarify the extent to which automated scores approximate genuine subjective experience. A related constraint is that this pipeline may partially reflect linguistic surface features of the reports rather than the underlying mental content, an inherent constraint of any text-based assessment that equally applies to human raters (Grizzard et al., 2025); integrating paralinguistic features (e.g., prosody, speech rate) of the audio recordings as complementary inputs to LLMs could improve annotation of less linguistically transparent dimensions.

Regarding the EEG decoding component, the overall predictive power found for Valence was modest, an outcome that may partly reflect the relatively limited number of observations both within (10 trials) and across (22) participants. The stringent methodological controls implemented to ensure that models decoded actual phenomenological content rather than participant identity, including within-subject median binarization, within-subject scaling, and within-subject permutation testing, may have conservatively attenuated classification performance. Larger samples with more trials per participant should strengthen both within-subject and group-level decoding and could reveal generalizable neural signatures for dimensions that appeared idiosyncratic in the current dataset. High-density EEG or concurrent fMRI recordings could also improve spatial resolution and access to medial structures implicated in self-referential and temporal processing (Kim et al., 2024).

In summary, the present study developed and applied a combined approach for the phenomenological exploration of spontaneous thought, integrating retrospective free reports, automated LLM-based annotation, and EEG decoding. The results indicate that current LLMs can automate phenomenological annotation at a level of agreement comparable to, or exceeding, inter-human consistency, while substantially reducing the time and cost associated with manual coding. The successful decoding of affective valence from resting-state EEG provides initial evidence that LLM-derived annotations carry biologically meaningful signal, though the modest effect size (AUC = 0.61) and the failure to decode other dimensions at the group level indicate that this approach remains a proof of concept. Future work with larger samples, higher-density recordings, and individualized modeling may clarify whether the neurophysiological signatures of non-affective phenomenological dimensions are absent, idiosyncratic, or simply below the detection threshold of the current design.

## 5 Code & Data Availability

Code and processed data are available at https://github.com/Nicobruno92/mw_free_speech. Raw data can be made available upon request.

## 6 Authors Contributions

Research design: N.B., F.C., F.Z., C.P. and E.T.; Data collection: N.B., F.C., F.Z., T.D., S.M. and L.A.F.; Data curation: N.B.; EEG analysis: N.B.; Machine learning models: N.B.; Writing—original draft: N.B., C.P. and E.T.; Writing—reviewing and editing: all authors; Project administration: C.P., E.T. and M.V.;

## 8 Supplemental material

### 8.1 Supplementary Methods

#### 8.1.1 Zero-Shot Prompt Configuration

The computational analysis of the verbal reports was conducted using a zero-shot prompting strategy. The following prompt encompasses the system instructions, the detailed phenomenological definitions for all 11 dimensions, the rating scale anchors, and the mandatory JSON output format. This configuration was maintained consistently across all evaluated LLMs.

**Figure S1:**
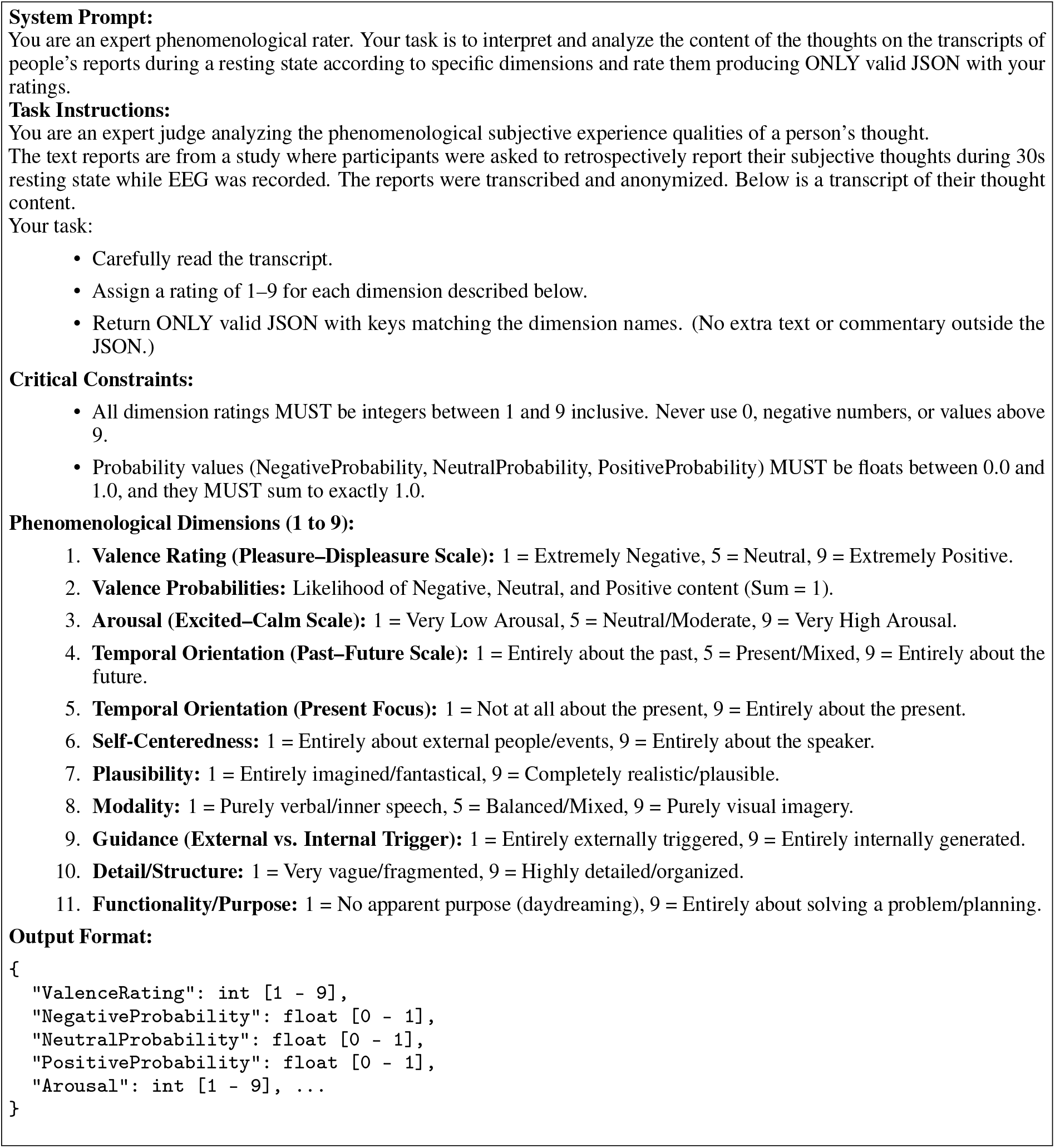
Zero-shot prompt configuration. Technical layout of the instructions and dimensions provided to the LLMs for automated phenomenological rating.

#### 8.1.2 Prompting Strategy

The influence of the sampling temperature, which regulates the degree of randomness in the model’s responses, was systematically examined. To evaluate stability and optimal performance, we compared a quasi-deterministic baseline (*T* = 0, single evaluation) against a stochastic aggregation approach at high temperature (*T* = 1). In the latter condition, the model’s rating procedure was repeated up to 10 independent times per report, allowing for greater diversity in generation, and the resulting ratings were subsequently averaged.

This approach allowed us to assess how the stability of the models’ ratings and their convergence toward the human gold standard (measured via Spearman’s rank correlation) evolved with repeated sampling. As illustrated in Supplementary Figure S2, while the stochastic ratings from a single high-temperature evaluation initially underperformed the deterministic baseline, aggregating multiple high-temperature evaluations rapidly improved performance across all models. The correlation plateaued after approximately 4 to 6 iterations, consistently outperforming the deterministic zero-temperature single-shot baseline. This finding confirmed that stochastic averaging leverages the internal variability of the LLMs to produce a more robust and consensus-aligned final rating.

**Figure S2:**
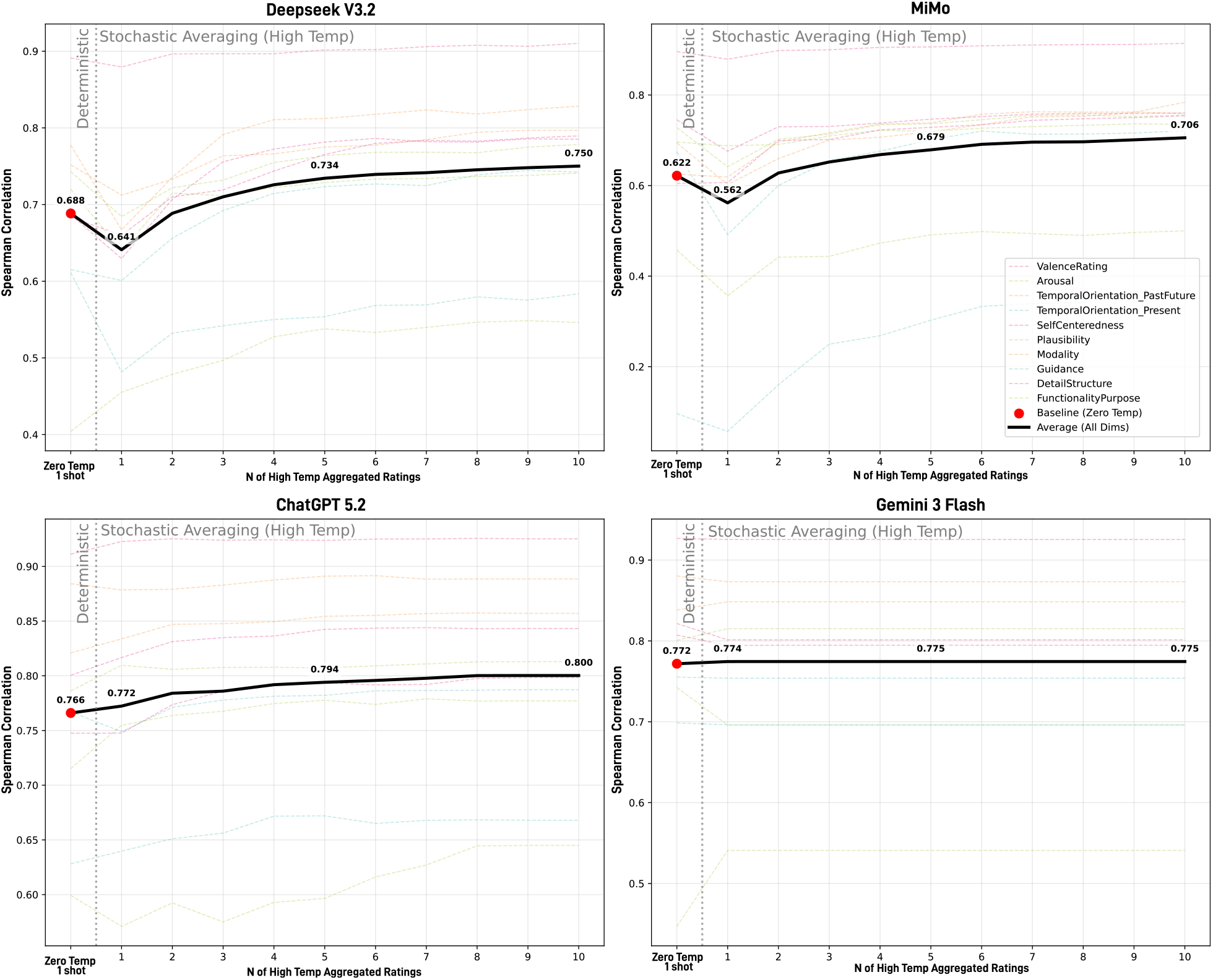
Effect of Stochastic Averaging on LLM-Human Agreement. The line plots illustrate how the Spearman’s rank correlation with human consensus evolves as a function of the number of aggregated high-temperature (*T* = 1) ratings, evaluated across four models (Deepseek V3.2, MiMo, ChatGPT-5.2, and Gemini-3-Flash). The solid black line represents the average correlation across all phenomenological dimensions, while dashed colored lines indicate individual dimensions. The red dot represents the baseline performance obtained from a single, quasi-deterministic zero-temperature (*T* = 0) evaluation. Across all models, aggregating multiple high-temperature ratings rapidly improved agreement with human consensus, ultimately outperforming the deterministic single-shot baseline and plateauing after approximately 4 to 6 iterations.

In addition to the baseline prompting, an alternative pseudo-Chain-of-Thought (CoT) strategy was evaluated under deterministic conditions (*T* = 0). This CoT approach instructed the model to follow a sequence of intermediate reasoning steps before delivering the final evaluation: first, to explicitly reason about and explain its interpretation of the specific dimension within the context of the report; second, to binarize the decision by determining the appropriate side of the semantic scale; and finally, to provide the continuous Likert rating. Although this reasoning-based strategy was designed to facilitate contextual understanding, comparative analysis utilizing DeepSeek-V3.2 indicated that the deterministic CoT approach yielded a lower overall agreement with human consensus (dimension-averaged *ρ* = 0.70) compared to the zero-shot stochastic averaging approach (dimension-averaged *ρ* = 0.75). The zero-shot high-temperature aggregation matched or outperformed the deterministic CoT strategy across the majority of individual phenomenological dimensions, with marked improvements observed in constructs such as Self-centeredness (*ρ* = 0.79 vs 0.71) and Detail/Structure (*ρ* = 0.79 vs 0.67). Based on these findings, the zero-shot stochastic averaging approach was retained as the primary methodology.

**Table S1:** Cost analysis of Large Language Models (LLMs) for automated phenomenological rating. Costs were estimated based on a single independent evaluation of 220 retrospective verbal reports (involving 220 API calls per run). Notably, prompt caching mechanisms were utilized for models supporting this functionality (e.g., ChatGPT and Gemini) to optimize token usage and reduce associated expenses.

| Model | Prompt Caching | Avg. Cost per Transcript (USD) | Total Cost per Run (USD) |
| --- | --- | --- | --- |
| DeepSeek-V3.2 | Yes | \$0.0005 | \$0.119 |
| MiMo | No | \$0.000 | \$0.00 |
| ChatGPT 5.2 | Yes | \$0.0046 | \$1.0611 |
| Gemini 3 Flash | Yes | \$0.0011 | \$0.2534 |

#### 8.1.3 EEG Markers

**Table S2:**
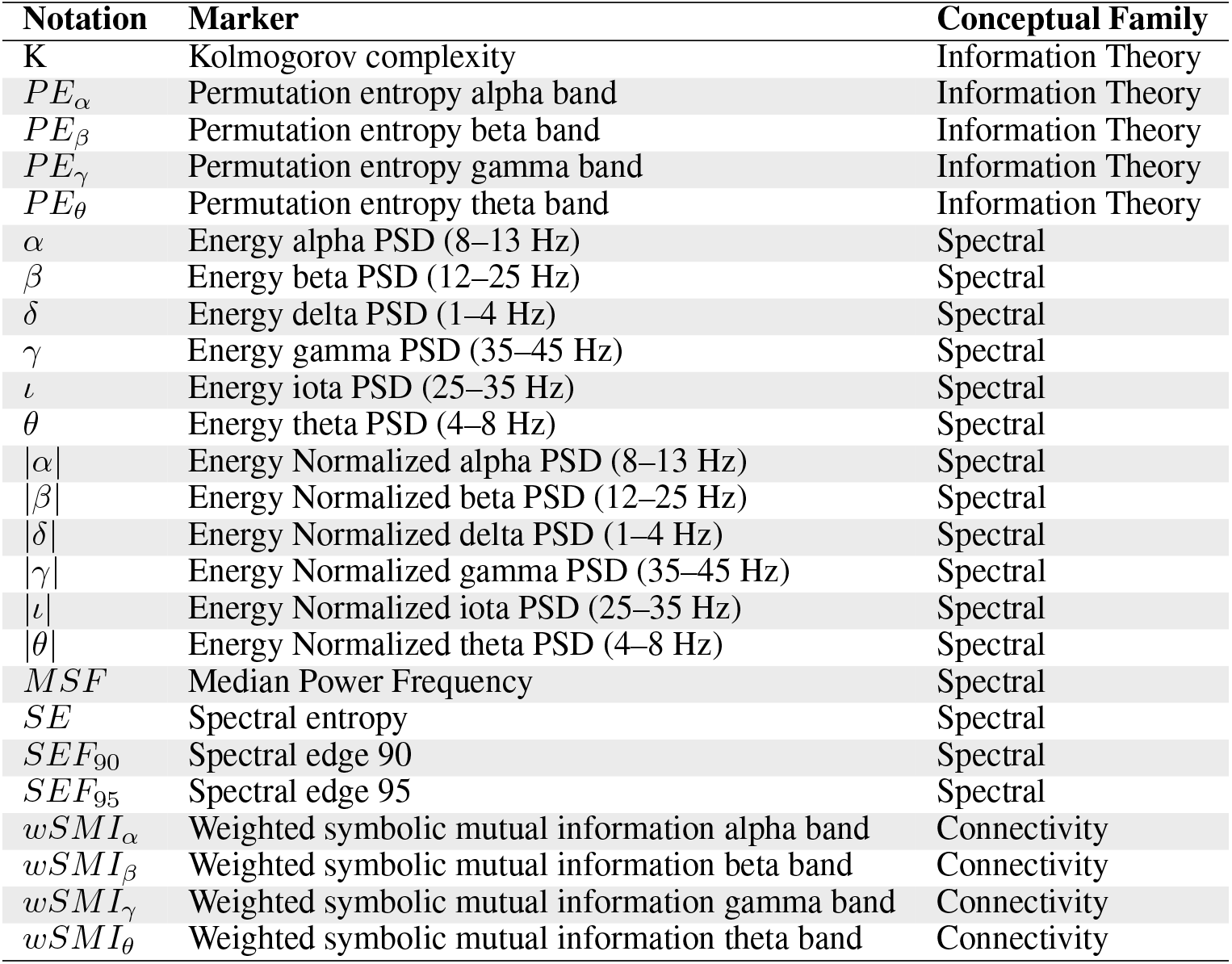
Overview of EEG markers employed in the present study, adapted from (Engemann et al., 2018; Sitt et al., 2014). The notation column presents the analytical symbols; the marker column details the complete description (including frequency bands for spectral measures); and the conceptual family denotes the theoretical categorization of each feature.

### 8.2 Supplemental Results

#### 8.2.1 Intraclass Correlation Coefficient Analysis

To complement the Spearman rank correlation analysis reported in the main text, ICC were computed between each LLM’s ratings and the human consensus across all ten phenomenological dimensions. While Spearman’s *ρ* captures the monotonic association between rankings, ICC provides a stricter measure of absolute agreement that accounts for both correlation and systematic bias in rating magnitude. As shown in Figure S3, the ICC values closely mirrored the Spearman correlations, confirming that LLMs not only preserved the rank ordering of human judgments but also approximated their absolute scale. ChatGPT-5.2 achieved the highest dimension-averaged ICC (0.79), followed by Gemini-3-Flash (0.78), DeepSeek-V3.2 (0.75), and MiMo (0.65), all exceeding the Human Rater Consistency baseline (ICC_human_ = 0.60). At the dimension level, the ICC pattern replicated the Spearman results: Valence exhibited the strongest absolute agreement (ICC = 0.90–0.92), followed by Past/Future (ICC = 0.85–0.91) and Goal Oriented (ICC = 0.77–0.89), while Arousal (ICC = 0.41–0.65) and Internally/Externally (ICC = 0.27–0.76) showed the most variability across models. The convergence between Spearman and ICC metrics indicates that the high LLM-human correlations reported in the main text reflect genuine agreement rather than an artifact of preserved rank order with divergent absolute values.

**Figure S3:**
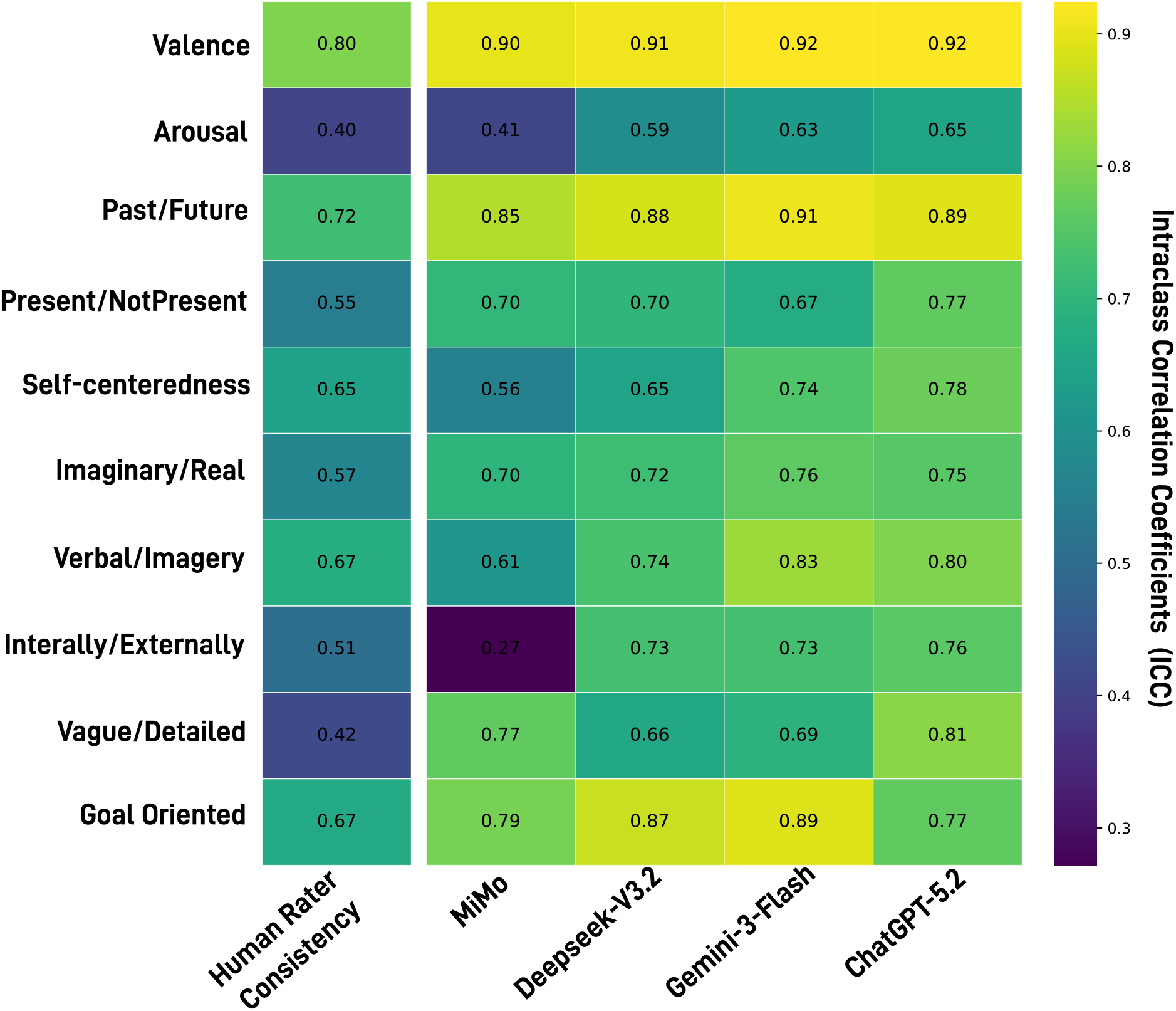
Intraclass Correlation Coefficients (ICC) between LLMs and Human Consensus across Phenomenological Dimensions. Heatmap displaying ICC values for four LLMs and the Human Rater Consistency baseline (Avg to Mean). The ICC metric captures absolute agreement, complementing the Spearman rank correlations reported in the main text (Figure 2). The pattern closely mirrors the Spearman results, with all LLMs exceeding the human baseline and the strongest agreement observed for Valence and Past/Future.

#### 8.2.2 Inter-Model Agreement Analysis

To further characterize the consistency of phenomenological ratings across LLM architectures, pairwise Spearman rank correlations were computed between all four models for each of the ten dimensions independently of their agreement with human consensus (Figure S4).

Inter-model agreement was high across most dimensions. Valence exhibited the strongest cross-architecture consensus (*ρ* = 0.92–0.96), followed by Goal Oriented (*ρ* = 0.82–0.90), Vague/Detailed (*ρ* = 0.76–0.90), and Verbal/Imagery (*ρ* = 0.76–0.90). Past/Future also showed robust agreement (*ρ* = 0.81–0.91), with the highest consistency between Gemini-3-Flash and ChatGPT-5.2 (*ρ* = 0.91). Self-centeredness (*ρ* = 0.75–0.88), Arousal (*ρ* = 0.79–0.89), and Present/NotPresent (*ρ* = 0.69–0.84) exhibited somewhat wider ranges, though still indicative of substantial agreement. The largest inter-model discrepancies were observed for Imaginary/Real (*ρ* = 0.57–0.83), where the DeepSeek-V3.2 and ChatGPT-5.2 pair reached *ρ* = 0.83 while MiMo showed markedly lower agreement with the remaining models, and for Internally/Externally (*ρ* = 0.46–0.75), the only dimension where some model pairs fell below *ρ* = 0.60.

These two dimensions also displayed the lowest LLM-human agreement and human inter-rater consistency in the main analysis (Figure 2), suggesting they reflect genuinely ambiguous constructs rather than architecture-specific interpretation failures.

**Figure S4:**
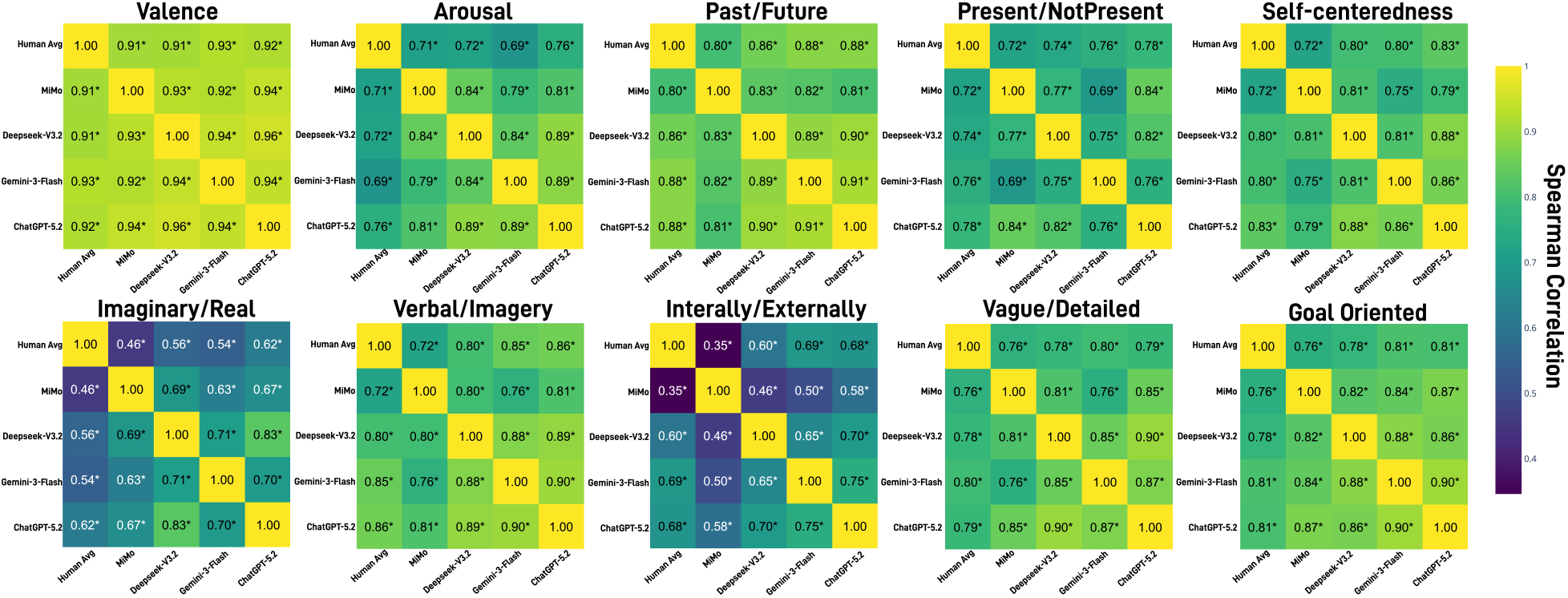
Inter-Model Agreement across Phenomenological Dimensions. Heatmaps displaying Spearman rank correlations between all LLM pairs (MiMo, DeepSeek-V3.2, Gemini-3-Flash, ChatGPT-5.2) for each of the ten phenomenological dimensions. Each panel represents a single dimension, with correlation coefficients indicating the degree of agreement between model pairs. Warmer colors reflect stronger inter-model consensus. Results demonstrate substantial cross-architecture agreement for most dimensions, particularly Valence, Past/Future, and Goal Oriented, indicating that different LLMs converge on similar interpretations of phenomenological constructs.

#### 8.2.3 Sentiment Analysis of Valence Ratings

To independently validate the high agreement observed between human raters and LLMs on the Valence dimension, an auxiliary comparative analysis was conducted using established sentiment analysis tools. Text descriptions from the retrospective free reports were processed using two well-known sentiment classifiers: VADER (Hutto & Gilbert, 2014), a lexicon and rule-based sentiment analysis tool specifically attuned to sentiments expressed in social media and short texts, and a RoBERTa-based model fine-tuned on ∼58 million tweets (Loureiro et al., 2022), representing a robust transformer-based approach to sentiment classification.

Spearman’s rank correlations were computed between the Valence ratings generated by ChatGPT-5.2 and the continuous sentiment scores produced by both VADER (compound score) and the RoBERTa model. As illustrated in Supplementary Figure S5, the analysis revealed substantial positive associations between the zero-shot LLM phenomenological ratings and both dedicated sentiment models. Specifically, ChatGPT-5.2 Valence scores exhibited a strong correlation with VADER sentiment scores (*ρ* = 0.72, *p <* 0.001) and an even stronger alignment with the RoBERTa predictions (*ρ* = 0.81, *p <* 0.001).

Importantly, the generative LLM demonstrated stronger agreement with the human consensus than both VADER and RoBERTa, which represent previous state-of-the-art approaches for this task. Furthermore, the LLM exhibited the highest inter-model agreement, effectively acting as a central hub that correlated strongly with both of these established tools. These findings suggest that the LLM, despite using a general zero-shot prompt based on the MDES scale, extracted emotional tone more reliably than models explicitly trained or designed for sentiment analysis. This independent validation further substantiates the robustness and efficacy of using advanced LLMs as automated raters for the affective dimensions of spontaneous thought.

**Figure S5:**
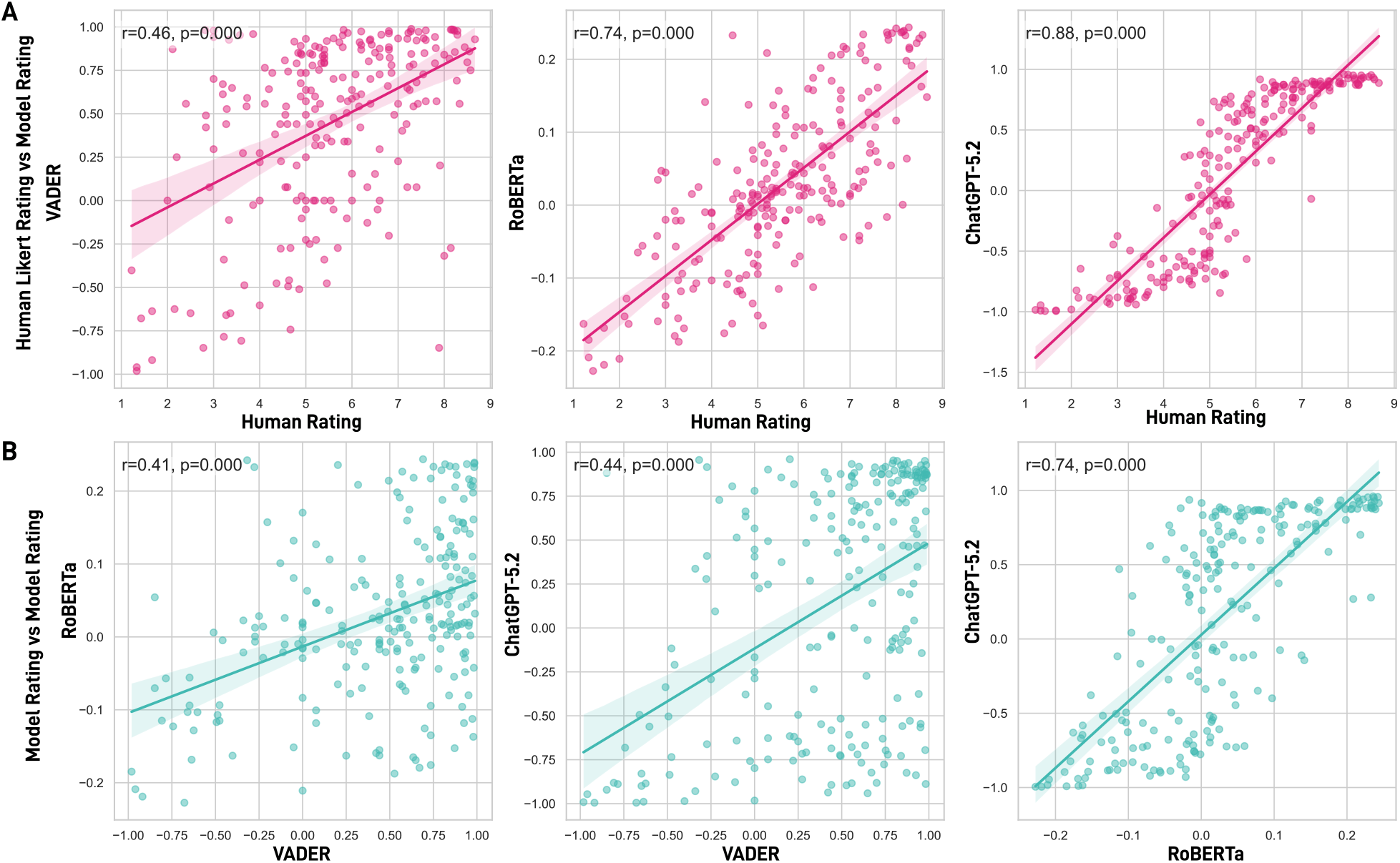
Comparison of ChatGPT-5.2 Valence Ratings with Established Sentiment Analysis Models. Scatter plots illustrating the association between the phenomenological Valence scores derived from ChatGPT-5.2 and continuous sentiment polarity scores generated by VADER (left) and a RoBERTa-based sentiment model (right) across all retrospective free reports. Each point represents a single report. Both comparisons yielded strong, statistically significant Spearman rank correlations (VADER: *ρ* = 0.72, *p <* 0.001; RoBERTa: *ρ* = 0.81, *p <* 0.001), independently validating the LLM’s capacity to accurately quantify the emotional tone of spontaneous thought.

#### 8.2.4 EEG Decoding of Sentiment Categories

Building upon the convergence between LLM-derived Valence ratings and established sentiment analysis tools (Section 8.2.3), a complementary decoding analysis was conducted to further characterize the neural discriminability of affective content. In addition to rating each report on a continuous Valence scale, ChatGPT-5.2 also provided probability estimates for three discrete sentiment categories, Positive, Neutral, and Negative, for each retrospective verbal report. These probability scores were binarized via within-subject median split and submitted to the same classification pipeline employed for the phenomenological dimensions (see 2).

Results indicated that the binary presence of Positive and Neutral sentiments (i.e., high vs. low probability) could be decoded from EEG features above chance level (Positive: AUC = 0.60, *p* = 0.02, FDR-corrected *p* = 0.03; Neutral: AUC = 0.58, *p* = 0.02, FDR-corrected *p* = 0.03; Supplementary Figure S6). The decoding of Negative sentiment exhibited a trend in the same direction but did not reach statistical significance (AUC = 0.55, *p* = 0.10, FDR-corrected *p* = 0.10). These findings corroborate and extend the main Valence result, indicating that the EEG signal contains discriminative information capable of differentiating moments characterized by high versus low positive and neutral affective tone, rather than exclusively reflecting a broad dimensional valence score.

**Figure S6:**
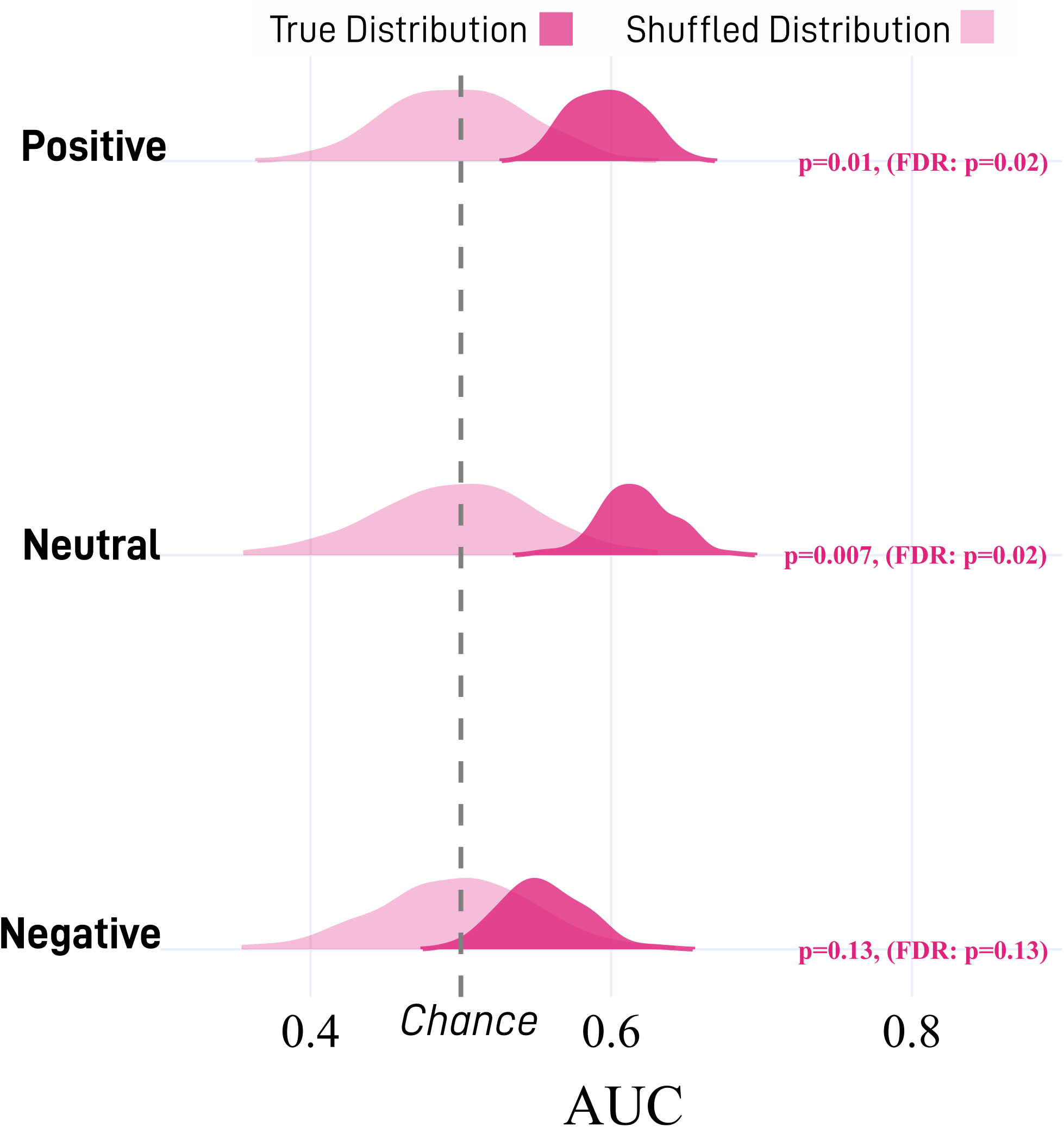
EEG Decoding of Sentiment Categories. Violin plots displaying the report-level AUC for the classification of Positive, Neutral, and Negative sentiment probabilities derived from ChatGPT-5.2, after averaging epoch-level predictions. Dark pink distributions represent true model AUC; light pink distributions represent the permutation-derived null. Positive and Neutral categories achieved statistically significant decoding (*p* = 0.02, FDR-corrected *p* = 0.03), while Negative did not reach significance (*p* = 0.10).

#### 8.2.5 Type 1 Error Validation Analysis

To ensure that the decoding models were not inadvertently learning to predict participant identity instead of the true phenomenological targets, we conducted a rigorous validation analysis using synthetic data. Surrogate datasets were generated by preserving each participant’s covariance structure while simultaneously destroying any genuine relationship to the phenomenological labels. The entire classification and evaluation pipeline—including epoch-level averaging and LOSO cross-validation—was subsequently applied to this null data.

We verified that the full pipeline maintained a Type 1 error rate below the *α* = 0.05 threshold across all phenomenological dimensions (Supplementary Figure S7). This signifies that false positive classification results (where the model trained on synthetic data spuriously outperformed the permutation distribution) occurred in fewer than 5% of the simulation runs. This validation step confirmed that the decoding pipeline was robust against spurious associations driven by subject-level data clustering, ensuring that the significant findings reported in the main text are attributable to genuine neuro-phenomenological relationships.

**Figure S7:**
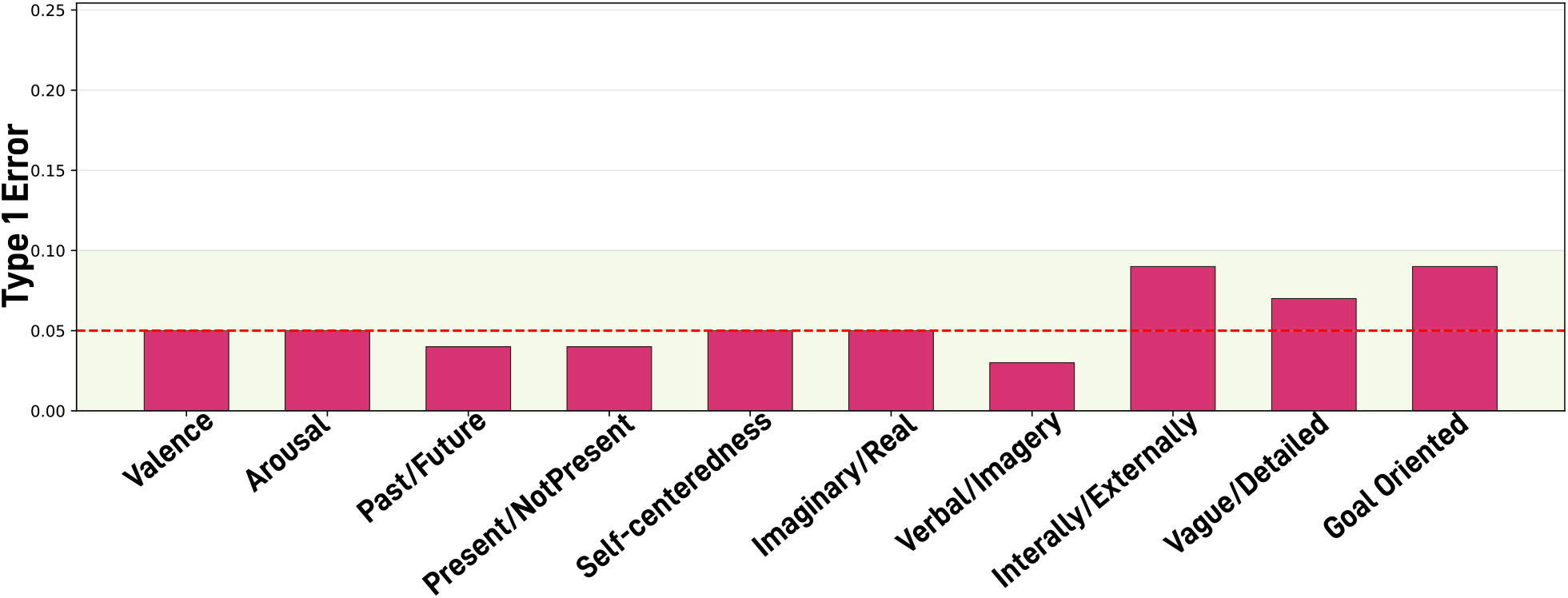
Type 1 Error Rates for EEG Decoding Models. Bar plot illustrating the Type 1 error rate across the ten phenomenological dimensions when the full classification pipeline was applied to synthetic null datasets. The red dashed line denotes the standard statistical significance threshold (*α* = 0.05). The green shaded area represents the acceptable 95% confidence interval for the error rate given the number of simulations. All dimensions demonstrated error rates consistent with or below the nominal alpha level, confirming that the pipeline does not systematically inflate false positives due to subject-level data structure.

#### 8.2.6 Feature Importance Across All Dimensions

To provide a comprehensive overview of the neurophysiological features driving the decoding models, Supplementary Figure S8 displays the full feature importance profiles across all ten phenomenological dimensions. While only Valence achieved group-level statistical significance in decoding performance, the exploratory patterns observed for the remaining dimensions may offer insights into their highly individualized neural correlates.

**Figure S8:**
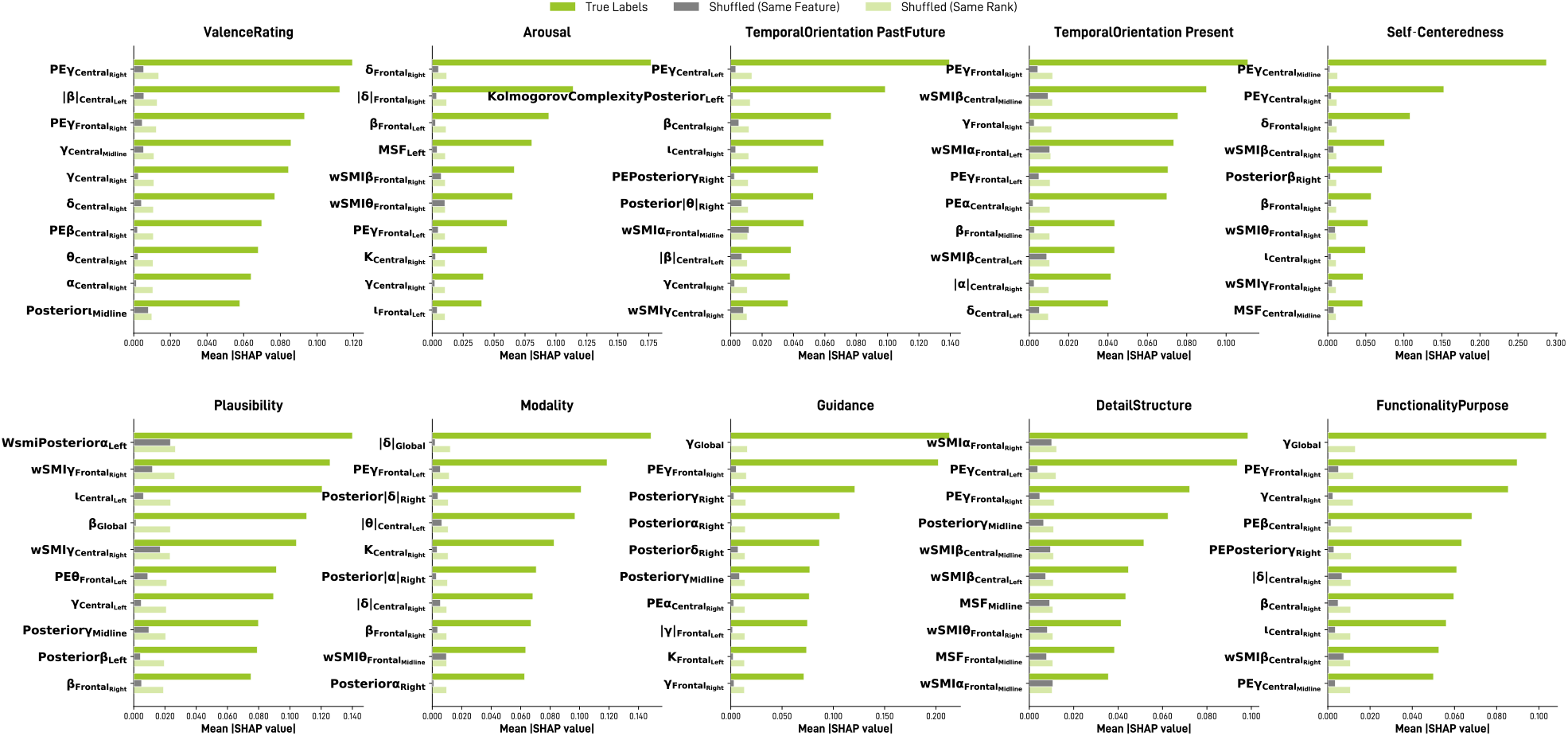
Feature Importance Profiles for All Phenomenological Decoding Models. Bar plots displaying the mean absolute SHAP values for the most discriminative EEG features across the ten phenomenological dimensions. The solid green bars (“True Labels”) represent the feature importance in the actual models, aggregated across all runs. To establish significance, these values are compared against two empirical baselines derived from permutation testing: the mean SHAP value of that exact same feature when labels were shuffled (dark grey, “Shuffled (Same Feature)”), and the mean SHAP value of whichever feature occupied that same rank position across shuffled runs (light green, “Shuffled (Same Rank)”). Features that substantially exceed both baselines reflect robust neurophysiological signatures.

